# A data compendium of *Mycobacterium tuberculosis* antibiotic resistance

**DOI:** 10.1101/2021.09.14.460274

**Authors:** Alice Brankin, Kerri M Malone, The CRyPTIC Consortium

## Abstract

The Comprehensive Resistance Prediction for Tuberculosis: an International Consortium (CRyPTIC) presents here a compendium of 15,211 *Mycobacterium tuberculosis* global clinical isolates, all of which have undergone whole genome sequencing (WGS) and have had their minimum inhibitory concentrations to 13 antitubercular drugs measured in a single assay. It is the largest matched phenotypic and genotypic dataset for *M. tuberculosis* to date. Here, we provide a summary detailing the breadth of data collected, along with a description of how the isolates were collected and uniformly processed in CRyPTIC partner laboratories across 23 countries. The compendium contains 6,814 isolates resistant to at least one drug, including 2,129 samples that fully satisfy the clinical definitions of rifampicin resistant (RR), multi-drug resistant (MDR), pre-extensively drug resistant (pre-XDR) or extensively drug resistant (XDR). Accurate prediction of resistance status (sensitive/resistant) to eight antitubercular drugs by using a genetic mutation catalogue is presented along with the presence of suspected resistance-conferring mutations for isolates resistant to the newly introduced drugs bedaquiline, clofazimine, delamanid and linezolid. Finally, a case study of rifampicin mono-resistance demonstrates how this compendium could be used to advance our genetic understanding of rare resistance phenotypes. The compendium is fully open-source and it is hoped that the dataset will facilitate and inspire future research for years to come.

## Introduction

Tuberculosis (TB) is a curable and preventable disease; 85% of those afflicted can be successfully treated with a six-month regimen. Despite this, TB is the world’s top infectious disease killer (current SARS-CoV-2 pandemic excepted) with 10 million new cases and 1.2 million deaths estimated in 2019 alone (1). Furthermore, drug resistant TB (DR-TB) is a continual threat; almost half a million cases resistant to the first-line drug rifampicin (RR-TB) were estimated, with three quarters of these estimated to be <u>m</u>ulti<u>d</u>rug-<u>r</u>esistant (MDR-TB, resistant to first-line drugs isoniazid and rifampicin) (1). Worryingly, only 44% of DR-TB cases were officially notified and just over half of these cases were successfully treated (57%) (1).

To address these issues, the World Health Organisation (WHO) is encouraging the development of better, faster and more targeted diagnostic and treatment strategies through its EndTB campaign (1). Of particular interest is universal drug susceptibility testing (DST). Conventionally, DST relies on lengthy (4 weeks minimum) culture-based methods that require strict biosafety conditions for *Mycobacterium tuberculosis.* The development of rapid genetics-based assays has decreased diagnostic time to as little as 2 hours through the detection of specific resistance conferring mutations *e.g.* the Cepheid Xpert® MTB/RIF test (2, 3). However, assay bias towards specific genic regions can result in misdiagnosis of resistance, the prescription of ineffective treatment regimens and subsequent spread of multi-drug resistant disease, as seen during an MDR outbreak in Eswatini (4–6). Furthermore, detection of rifampicin resistance is used to infer MDR-TB epidemiologically as rifampicin resistance tends to coincide with resistance to isoniazid (7). While this *modus operandi* is successful at pragmatically identifying potential MDR cases quickly and effectively, it is not generally true that a single path exists for developing MDR or e<u>x</u>tensively <u>d</u>rug <u>r</u>esistant TB (XDR = MDR/RR + resistance to at least one fluoroquinolone and either bedaquiline or linezolid).

Whole-genome sequencing (WGS) has the potential to reveal the entirety of the *M. tuberculosis* genetic resistance landscape for any number of drugs simultaneously whilst enabling a more rapid turnaround time and reduction in cost compared to DST culture-based methods (8). However, the success of WGS as a diagnostic tool wholly depends on there being a comprehensive and accurate catalogue of resistance-conferring mutations for each drug. Recent advances have shown that genotypic predictions of resistance correlate well with DST measurements for first-line drugs (7). However, the mechanisms of resistance to second-line drugs along with the new and re-purposed drugs (NRDs) are less well understood despite their increased administration in clinics as MDR cases climb(1, 9).

To address these shortcomings, the Comprehensive Resistance Prediction for Tuberculosis: an International Consortium (CRyPTIC) has collected *M. tuberculosis* clinical isolates worldwide to survey the genetic variation associated with resistance to 13 antitubercular drugs, specifically the first-line drugs rifampicin, isoniazid, ethambutol, the second-line drugs amikacin, kanamycin, rifabutin, levofloxacin, moxifloxacin, ethionamide, and the new and re-purposed drugs bedaquiline, clofazimine, delamanid and linezolid. Here, we introduce and describe these data in the form of an open-access compendium of 15,211 isolates, each of which has had its genomic sequence determined and DST profile measured (10). This compendium is the largest drug screening effort to date for *M. tuberculosis* in a ‘one isolate – one microscale assay’ format across defined compound concentration ranges. The presented dataset forms the backbone for several studies being put forth by the consortium to achieve its ultimate aim (11–15). By being fully open-access, it is hoped that this compendium will prove an invaluable resource to accelerate and improve antimicrobial resistance (AMR) diagnostic development for TB, both by the enrichment of mutation catalogues for WGS resistance prediction and the identification of important diagnostic gaps and drug resistance patterns.

## Methods

### Ethics

Approval for the CRyPTIC study was obtained by Taiwan Centers for Disease Control IRB No. 106209, University of KwaZulu Natal Biomedical Research Ethics Committee (UKZN BREC) (reference BE022/13) and University of Liverpool Central University Research Ethics Committees (reference 2286), Institutional Research Ethics Committee (IREC) of The Foundation for Medical Research, Mumbai (Ref nos. FMR/IEC/TB/01a/2015 and FMR/IEC/TB/01b/2015), Institutional Review Board of P.D. Hinduja Hospital and Medical Research Centre, Mumbai (Ref no. 915-15-CR [MRC]), scientific committee of the Adolfo Lutz Institute (CTC-IAL 47-J / 2017) and in the Ethics Committee (CAAE: 81452517.1.0000.0059) and Ethics Committee review by Universidad Peruana Cayetano Heredia (Lima, Peru) and LSHTM (London, UK).

### Sample collection

Participating collection centres varied in their isolate collection approaches and timescales *(e.g.* longitudinal sampling, rolling patient visits, biobank stocks), but the consortium collectively aimed to oversample for *M. tuberculosis* isolates with drug resistance and multi-drug resistance. A standard operating protocol for sample processing was defined by CRyPTIC as previously described and is discussed in more detail below in relevant sub-sections (10, 11).

### Plate assay

The CRyPTIC consortium designed two versions of the Sensititre MYCOTB plate (Thermo Fisher Scientific Inc., USA) named the “UKMYC5” and “UKMYC6” microtitre plates (10, 11). These plates contain five to ten doubling dilutions of 13 antibiotics (rifampicin (RIF), rifabutin (RFB), isoniazid (INH), ethambutol (EMB), levofloxacin (LEV), moxifloxacin (MXF), amikacin (AMI), kanamycin (KAN), ethionamide (ETH), clofazimine (CFZ), linezolid (LZD), delamanid (DLM), and bedaquiline (BDQ)). Delamanid and bedaquiline were provided by Otsuka Pharmaceutical Co., Ltd. and Janssen Pharmaceutica respectively. The UKMYC5 plate also contained para-aminosalicylic acid (PAS), but the MICs were not reproducible and hence it was excluded from the UKMYC6 plate design and is not included in any subsequent analysis (10).

A standard operating protocol for sample processing was defined by CRyPTIC as previously described (10, 11). Clinical samples were sub-cultured using 7H10 agar plates, Lowenstein-Jensen tubes or MGIT tubes. Bacterial cell suspensions (0.5 McFarland standard, saline Tween) prepared from (no later than) 14-day old colonies were diluted 100X in 10 ml enriched 7H9 broth prior to plate inoculation. A semi-automated Sensititre Autoinoculator (Thermo Fisher, Scientific Inc., USA) was used to inoculate 100 µl prepared cell suspensions (1.5 x 10^5^ CFU/ml [5 x 10^4^ CFU/ml - 5 x 10^5^ CFU/ml]) into each well of a UKMYC5/6 microdilution plate. The plate was sealed and incubated for 14 days at 37°C. Quality control runs were performed periodically using *M. tuberculosis* H37Rv ATCC 27294, which is sensitive to all drugs on the plates.

### Minimum Inhibitory Concentration (MIC) measurements

Minimum inhibitory concentrations (MICs) for each drug were read after incubation for 14 days by a laboratory scientist using a Thermo Fisher Sensititre™ Vizion™ digital MIC viewing system (10). The Vizion apparatus was also used to take a high contrast photograph of the plate with a white background, from which the MIC was measured again using the Automated Mycobacterial Growth Detection Algorithm (AMyGDA) software (16). The AMyGDA algorithm was specifically developed to automate and perform quality control of MIC measurements, and to facilitate machine learning studies within the consortium. AMyGDA detects the boundaries of each well using a Hough transform for circles and measures growth as the number of dark pixels within the area contained by this boundary.

All images where the MICs measured by Vizion and AMyGDA were different were uploaded to a citizen science project, BashTheBug, on the Zooniverse platform (17). Each image was then classified by ≥11 volunteers and the median classification taken. MICs were then classified as high (at least two methods concur on the MIC), medium (either a scientist recorded a MIC measurement using Vizion but did not store the plate picture, or Vizion and AMyGDA disagree and there is no BashTheBug measurement), or low (all three methods disagree) quality.

To ensure adequate data coverage for *this* study, we took the MIC from the Vizion reading provided by the trained laboratory scientist if it was annotated as having medium or low quality.

### Binary phenotype classification

Binary phenotypes (resistant/susceptible) were assigned from the MICs by applying epidemiological cut-off values (11); samples with MICs at or below the ECOFF are, by definition, wild-type and hence assigned to be susceptible to the drug in question (11). Samples with MICs above the ECOFF are therefore classified as resistant (Fig. S1, Table S1). Please see (11) for the body of work supporting the use of the ECOFF relative to the compendium isolates and supplemental Table S1 for the ECOFFs for each drug tested.

### Genomic data processing and variant calling

Short-read, paired end libraries were sequenced on Illumina machines and the resulting FASTQ files were processed using the bespoke pipeline Clockwork (v0.8.3, github.com/iqbal-lab-org/clockwork, (18)). Briefly, all raw sequencing files were indexed into a relational database with which Clockwork proceeds. Human, nasopharyngeal flora and human immunodeficiency virus related reads were removed and remaining reads were trimmed (adapters and low quality ends) using Trimmomatic and mapped with BWA-MEM to the *M. tuberculosis* H37Rv reference genome (NC000962.3) (19, 20). Read duplicates were removed. Genetic variants were called independently using Cortex and SAMtools, two variant callers with orthogonal strengths (SAMtools a high sensitivity SNP caller, and Cortex a high specificity SNP and indel caller) (21, 22). The two call sets were merged to produce a final call set, using the Minos adjudication tool to resolve loci where the two callers disagreed, by remapping reads to an augmented genome containing each alternative allele (23). Default filters of a minimum depth of 5x, a fraction of supporting reads of 0.9 (minos) and a genotype confidence percentile (GCP) filter of 0.5 were applied. The GCP filter is a normalised likelihood ratio test, giving a measure of confidence in the called allele compared with the other alternatives, and is described in (23). This produced one variant call format (VCF) file per sample, each only describing positions where that sample differed from the reference.

These filtered VCFs were then combined, to produce a single non-redundant list of all variants seen in the cohort. All samples were then processed a second time with Minos, remapping reads to a graphical representation of all the segregating variation within the cohort, generating VCF files which had an entry at all variable positions (thus for all samples, most positions would be genotyped as having the reference allele). These “regenotyped” VCFs were later used to calculate pairwise distances (see below).

To remove untrustworthy loci, a genome mask was applied to the resulting VCF files (regions identified with self-blast matches in (24) comprising of 324,971 bp of the reference genome). Furthermore, positions with less than 90% of total samples passing default Clockwork/Minos variant call filters (described above) were filtered out, comprising 95,703 bp of the genome, of which 55,980 bp intersect with the genome mask.

### Resistance prediction using a genetic catalogue

A hybrid catalogue of genetic variants associated with resistance to first- and second- line drugs based on existing catalogues was created and can be found at github.com/oxfordmmm/tuberculosis_amr_catalogues/blob/public/catalogues/NC_000962.3/NC_000962.3_CRyPTIC_v1.311_GARC1_RUS.csv (7, 25). We specifically did not use the recent WHO catalogue to avoid circularity and over-training, as that catalogue was developed (via prior literature, expert rules and a heuristic algorithm) based partially on these isolates (26)).

The resulting VCF file for each isolate (see “*Genomic data processing and variant calling”* section above) was compared to the genetic catalogue to determine the presence or absence of resistance-associated mutations for eight drugs: rifampicin, isoniazid, ethambutol, levofloxacin, moxifloxacin, amikacin, kanamycin and ethionamide. We did not apply the approach used in (7) to make a prediction if a novel mutation was detected in a known resistance gene, as we simply wanted to measure how well a pre-CRyPTIC catalogue could predict resistance in the compendium. These results (found in PREDICTIONS.csv, see “Data availability” section for access) were then compared to the binary phenotypes (see “Binary phenotype classification” section for how these were defined) with the following metrics calculated: TP: the number of phenotypically resistant samples are that correctly identified as resistant (“true positives”); FP, the number of phenotypically susceptible samples that are falsely identified as resistant (“false positives”); TN, the number of phenotypically susceptible samples that are correctly identified as susceptible (“true negatives”); FN, the number of phenotypically resistant samples that are incorrectly identified as susceptible (“false negative”); VME, very major error rate (false-negative rate), 0-1; ME, major error rate (false-positive rate), 0-1; PPV, positive predictive value, 0-1; NPV, negative predictive value.

### Phylogenetic tree construction

A pairwise genetic distance matrix was constructed for 15,211 isolates by comparing pairs of regenotyped VCF files (see “*Genomic data processing and variant calling”* section above for more details). A neighbourhood-joining tree was constructed from the distance matrix using *quicktree* (27). Tree visualisation and annotation was performed using the R library *ggtree* (28). *M. tuberculosis* lineages were assigned using Mykrobe and are represented by the coloured dots at the branch termini of the tree (23). For isolates that had ‘mixed’ lineage classification (*i.e.,* 2 lineages were found present in the sample by Mykrobe, *n* = 225, 1.5%), the first of the two lineages was assigned to the isolate. *ggtree* was also used to construct the trees depicting bedaquiline, clofazimine and delamanid resistant isolates.

### The Data

All data can be found at ftp.ebi.ac.uk/pub/databases/cryptic/reuse/. The FTP site contains two top level directories: “reuse” and “reproducibility”. In total, the compendium contains data entries for 15,211 isolates. Note that various filtering criteria have been applied in both this study and other CRyPTIC studies and thus final isolate numbers presented across the publications will vary.

#### “reuse” directory

We point the reader to this directory to gain access to CRyPTIC project data. “CRyPTIC_reuse_table_20221019.csv” contains genotypic and phenotypic data relating to the figures and summaries listed in this manuscript and is what we present as a general use reference table for most future projects. It includes binary phenotypes (R/S), MICs, phenotype quality metrics, and ENA sample IDs for 12,288 compendium isolates (see “Quality assurance of the minimum inhibitory concentrations for 13 drugs” section below in Results for filters applied to obtain this final set of isolated). It also includes file paths to each isolate’s VCF file and ‘regenotyped’ VCF file (VCF files which have an entry at all variable positions, see “Genomic data processing and variant calling” section above for more).

#### “reproducibility” directory

This directory contains the data used for multiple CRyPTIC project publications referenced throughout this manuscript. As stated above, each project has taken slightly different subsets of this data as documented in those papers. For example, see how tables such as “MUTATIONS.csv” and “GENOTYPES.csv” were used and filtered, (along with others) in this study to obtain the reuse file “CRyPTIC_reuse_table_20221019.csv” in Figure 1. Again, for optimal use of CRyPTIC data in your own project, please refer to “CRyPTIC_reuse_table_20221019.csv” in the “reuse” directory. All data for this study were analysed and visualised using either R or python3 libraries and packages. See github.com/kerrimalone/Brankin_Malone_2022 for codebase.

**Figure 1:**
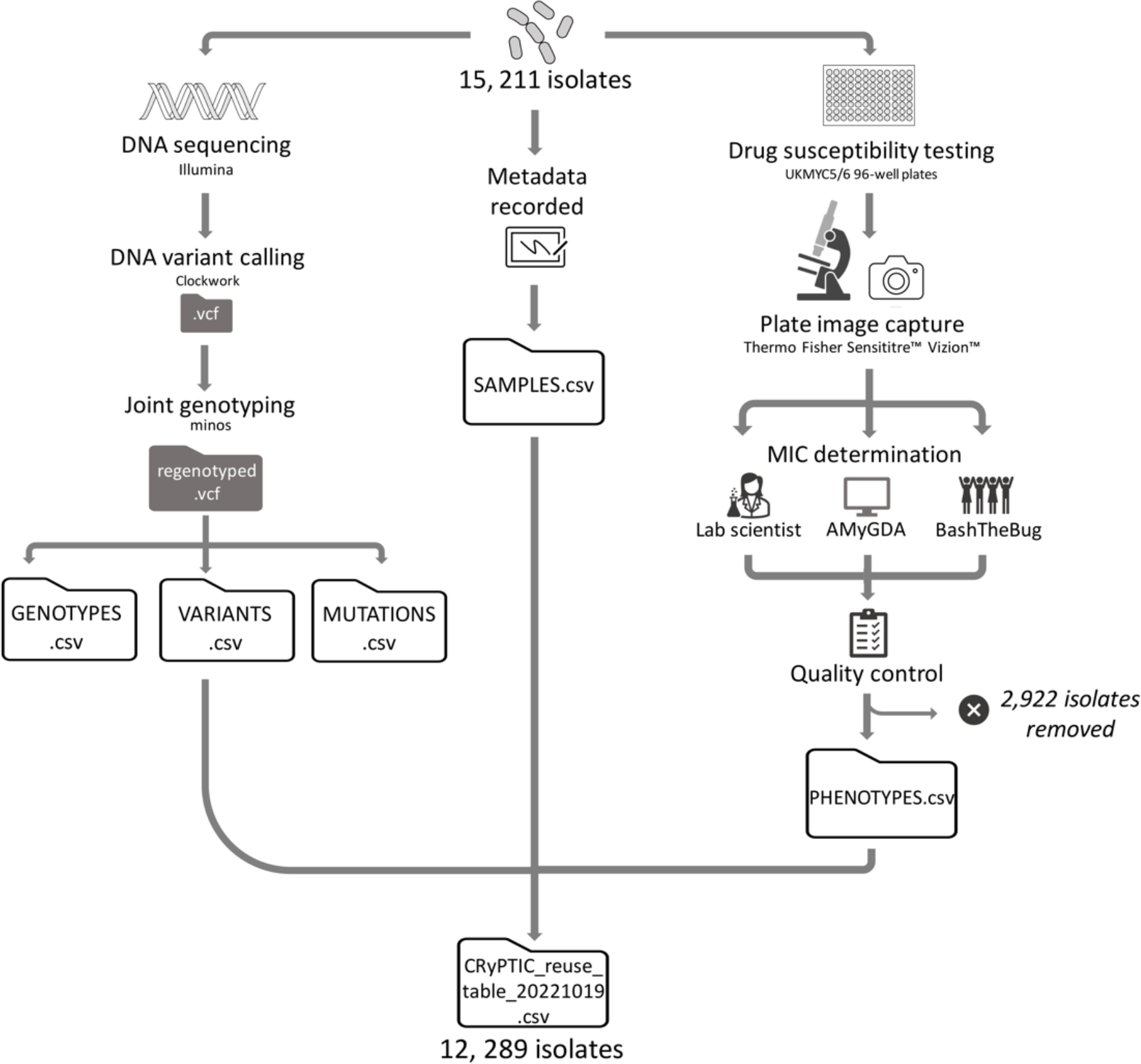
Processing sequencing and MIC data for 15,211 M. tuberculosis isolates. Briefly: Each isolate was DNA sequenced using an Illumina machine and plated onto 96 well plates (UKMYC5/6) containing 5-10x doubling dilutions of 13 antitubercular drugs for drug susceptibility testing. Associated metadata (including country of origin and processing laboratory) was recorded. DNA variant calling and analysis was performed using Clockwork and minos. After 14 days, MIC measurements were measured by a trained scientist using Vizion, and the plate was photographed to measure the MIC using the automated AMyGDA software and citizen scientists from BashTheBug. After quality control procedures, phenotypic MIC data for 2,922 isolates were removed. The compendium therefore contains 15,211 isolates with whole genome sequencing data, 12,289 of which have matched phenotypic data, presented in “CRyPTIC_reuse_table_20211019.csv” via an FTP site (see methods). The data tables GENOTYPES.csv, VARIANTS.csv, MUTATIONS.csv, SAMPLES.csv and PHENOTYPES.csv used for the analyses presented in this manuscript are also accessible via the FTP site (see methods).

## Results

### 15,211 M. tuberculosis clinical isolates

The CRyPTIC compendium contains 15,211 isolates for which both genomic and phenotypic data was collected by 23 of the partner countries across the continents of Asia, Africa, South America and Europe. An overview of the processing of the isolates is presented in Figure 1, and for a full description please see Methods.

The 15, 211 isolates originated from 23 different countries (Fig. 2). The largest number of isolates were contributed by India (*n =* 4,004), Peru (*n* = 2,691), South Africa (*n* = 2,155), Vietnam (*n* = 1,288) and China (*n* = 1,121).

**Figure 2:**
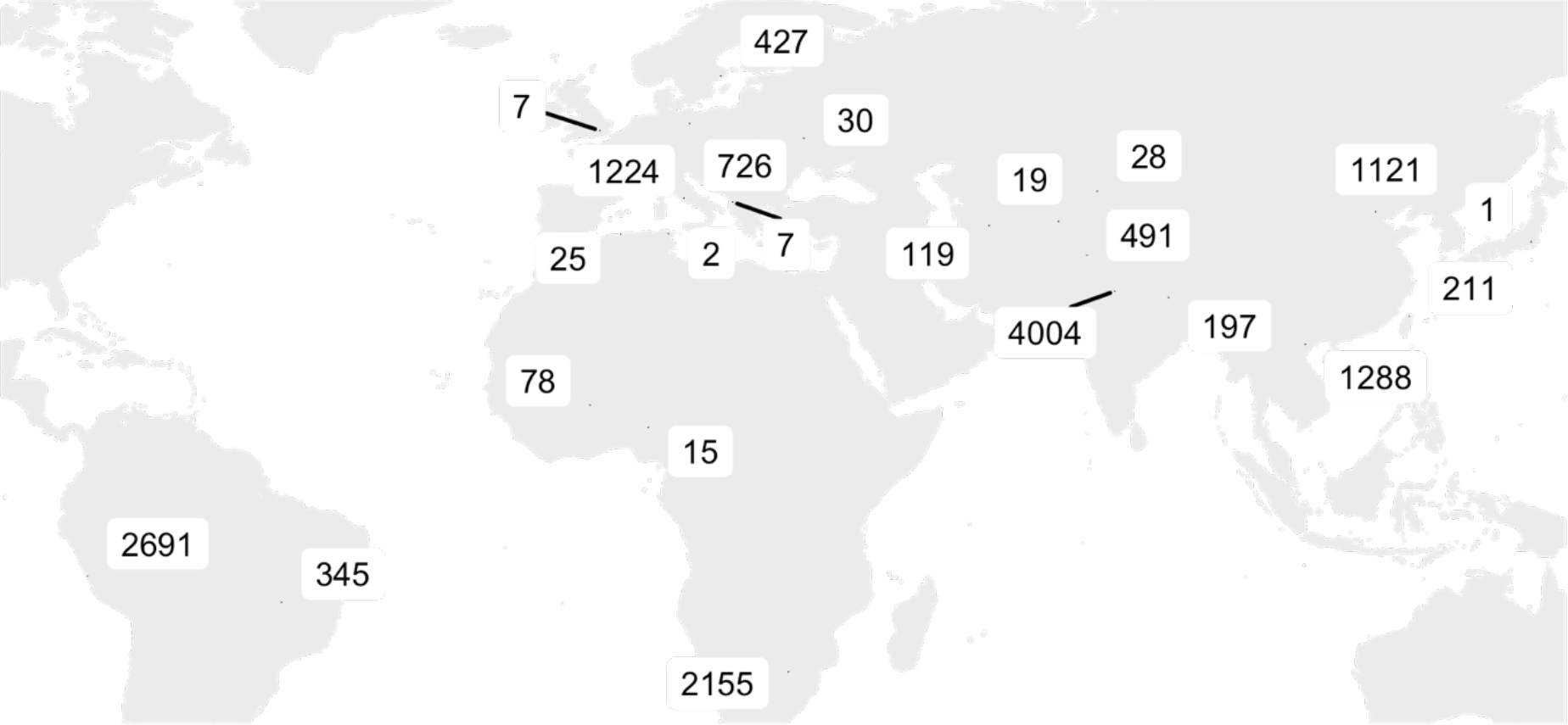
Geographical distribution of CRyPTIC M. tuberculosis clinical isolates. The total number of isolates contributed by each country is depicted. Where labels overlap, labels are exploded and lines are used to indicate country of origin, from left to right: UK, Albania, India. Where the origin of an isolate was not known, the collection site identity was assigned (this occurred for 269 isolates in Germany, 17 isolates in India, 6 isolates in Peru, 885 isolates in Italy, 510 isolates in South Africa, 357 isolates in Sweden, 208 isolates in Taiwan, 1 isolate in Brazil and 4 isolates in the UK).

Lineage assignment revealed that 99.7% of the 15,211 isolates belong to the four main *M. tuberculosis* lineages (L1-L4). The pie-charts in supplemental Figure S2 show the proportion of isolates amongst the different lineages (Table S2) and sub- lineages (Table S3) for each location. Like previous studies, we see a strong association between geolocation and lineage (Pearson’s chi-squared test, *p* < 2.2e- 16, Fig. S3) (29, 30). The phylogenetic tree in supplemental figure S4 further highlights the strong population structure of this collection of isolates, with isolates clustering according to lineage. Typically under-sampled in current databases and biobanks, the L3 isolates in this study represent the largest collection to date (31).

Although these 15,211 *M. tuberculosis* isolates were plated to determine their MICs to 13 antitubercular drugs, regular quality assurance checks detected problems with plate inoculation and reading in two laboratories. Therefore, 2,922 isolates were removed from the dataset, leaving a total of 12,289 isolates with matched phenotypic and genotypic data for further analysis (Fig.1). Due to wells being skipped and other phenomena that prevent an MIC being measured, 88.1% of the isolates had a phenotype for all 13 drugs on the plate. For each drug, the number of isolates with an MIC measurement, and the associated quality of the reading, is presented in Table 1.

**Table 1:**
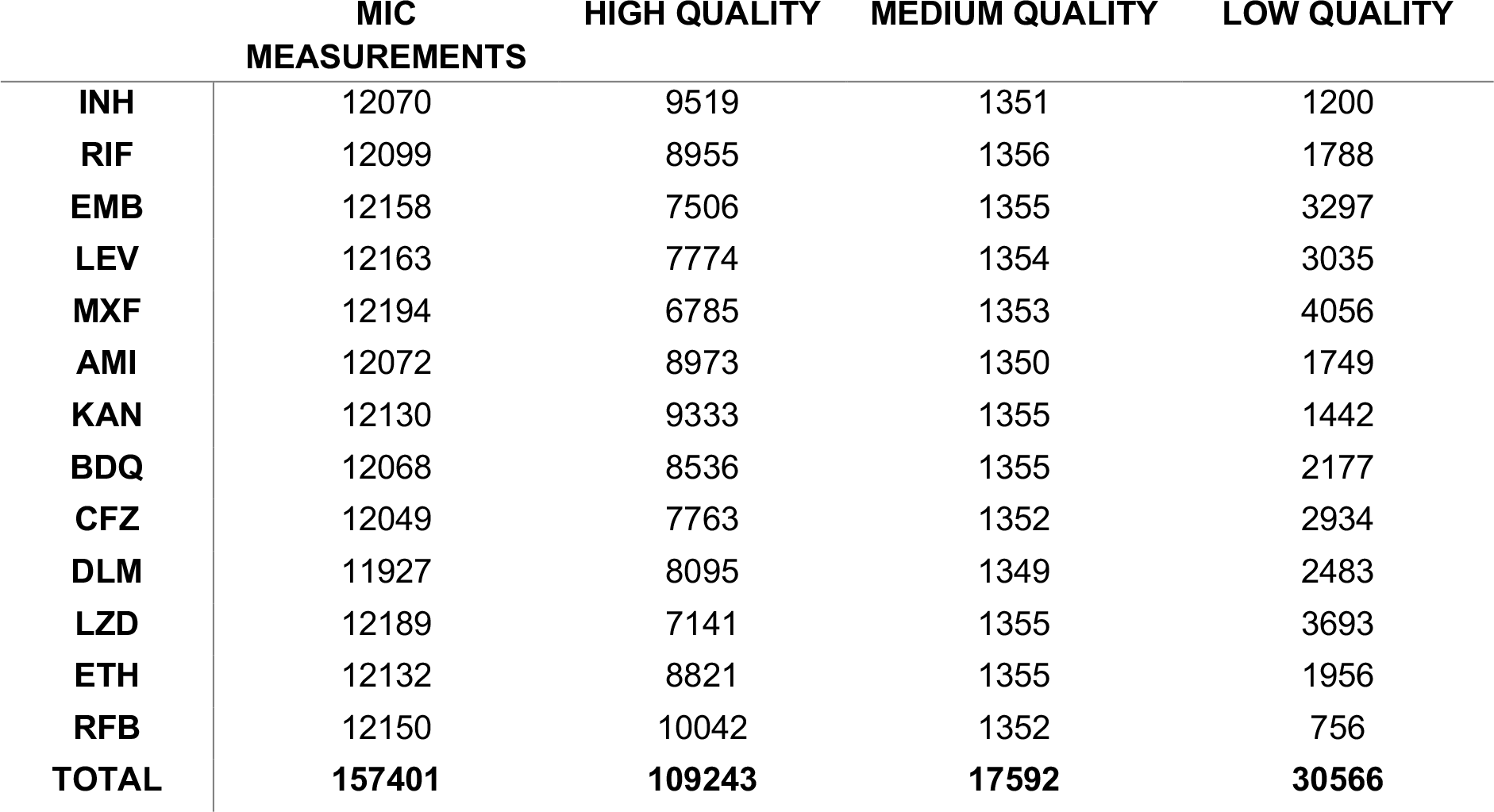
Quality metrics for phenotype data. Stated for each drug is the total number of MIC measurements stratified into “high” quality (at least two MIC measurement methods agree), “medium” quality (either Vizion and AMyGDA disagree, or the scientist recorded a MIC measurement using Vizion but did not store the plate picture) or “low” quality (all three MIC measurements methods disagree) phenotype classifications as described in Methods. Drug acronyms: INH = isoniazid, RIF = rifampicin, EMB = ethambutol, LEV = levofloxacin, MXF = moxifloxacin, AMI = amikacin, KAN = kanamycin, BDQ = bedaquiline, CFZ = clofazimine, DLM = delamanid, LZD = linezolid, ETH = ethionamide, RFB = rifabutin.

|  | MIC<br>MEASUREMENTS | HIGH QUALITY | MEDIUM QUALITY | LOW QUALITY |
| --- | --- | --- | --- | --- |
| INH | 12070 | 9519 | 1351 | 1200 |
| RIF | 12099 | 8955 | 1356 | 1788 |
| EMB | 12158 | 7506 | 1355 | 3297 |
| LEV | 12163 | 7774 | 1354 | 3035 |
| MXF | 12194 | 6785 | 1353 | 4056 |
| AMI | 12072 | 8973 | 1350 | 1749 |
| KAN | 12130 | 9333 | 1355 | 1442 |
| BDQ | 12068 | 8536 | 1355 | 2177 |
| CFZ | 12049 | 7763 | 1352 | 2934 |
| DLM | 11927 | 8095 | 1349 | 2483 |
| LZD | 12189 | 7141 | 1355 | 3693 |
| ETH | 12132 | 8821 | 1355 | 1956 |
| RFB | 12150 | 10042 | 1352 | 756 |
| TOTAL | 157401 | 109243 | 17592 | 30566 |

### Resistance classification and distribution

Unsurprisingly, given its size and bias toward the collection of resistant isolates, resistance to each of the 13 drugs is represented within the compendium (Fig. 3a). The drugs with the highest percentage of resistant isolates are the first line drugs isoniazid and rifampicin (49.0% and 38.7% respectively). Of the second line drugs, levofloxacin had the highest proportion of resistant isolates in the dataset (17.6%) and amikacin the lowest (7.3%). A low proportion of isolates were resistant to the NRDs, bedaquiline (0.9%), clofazimine (4.4%), delamanid (1.6%) and linezolid (1.3%).

**Figure 3:**
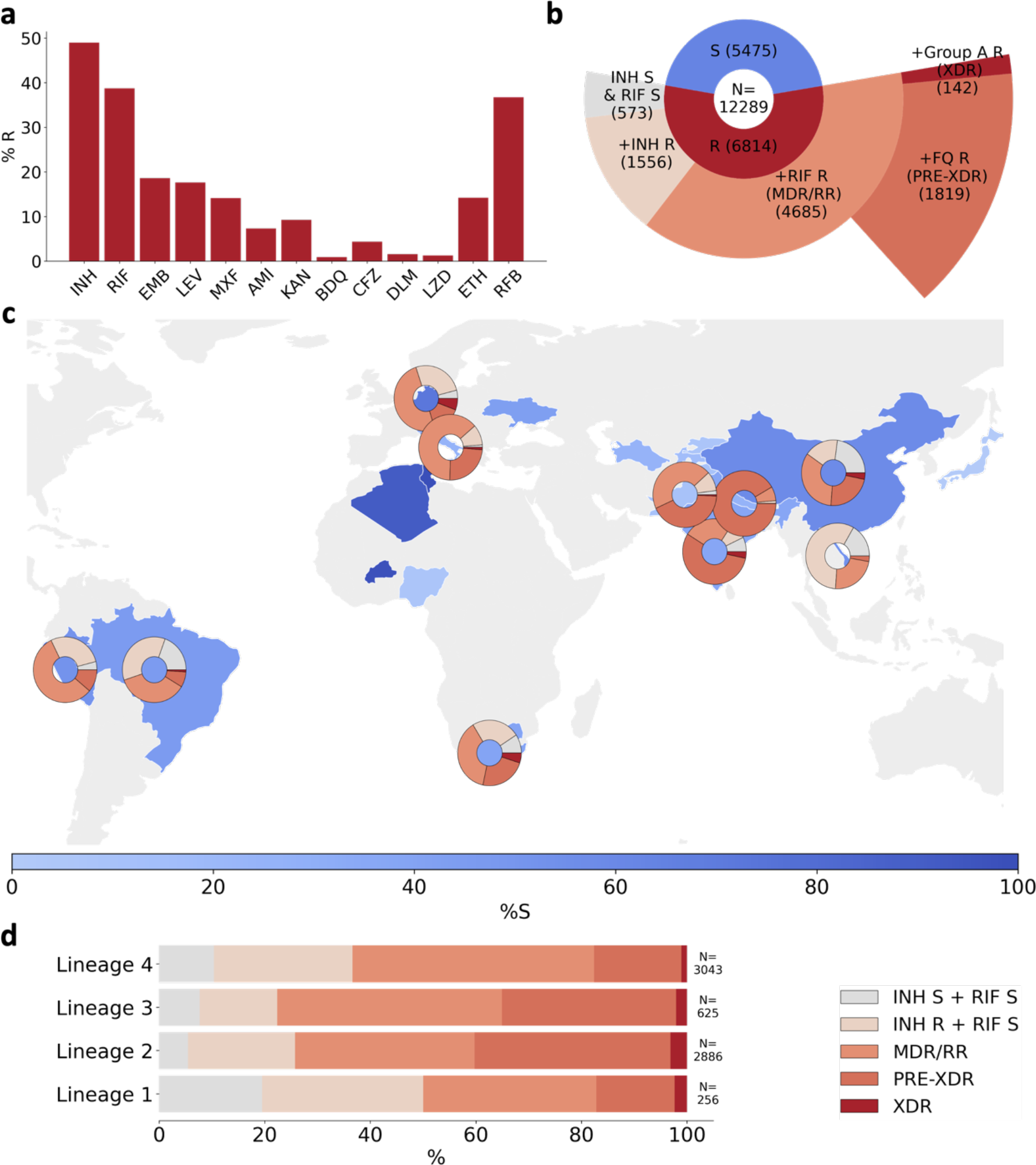
Drug phenotype data for the CRyPTIC compendium. **a)** Frequency of resistance to each of 13 drugs in the CRyPTIC dataset. The total number of isolates with a binary phenotype (of any quality) for the corresponding drug is presented in Table 1. **b)** Phenotypes of 12,289 CRyPTIC isolates with a binary phenotype for at least one drug. **c)** Geographical distribution of phenotypes of 12,289 CRyPTIC isolates. Intensity of blue shows the percentage of isolates contributed that were categorised as susceptible to all 13 drugs. Donut plots show the proportions of resistant phenotypes identified in (b) for countries contributing >=100 isolates with drug resistance. **d)** Proportions of resistance phenotypes in the 4 major *M. tuberculosis* lineages. N is the number of isolates of the lineage called resistant to at least one of the 13 drugs. Drug acronyms: INH = isoniazid, RIF = rifampicin, EMB = ethambutol, LEV = levofloxacin, MXF = moxifloxacin, AMI = amikacin, KAN = kanamycin, BDQ = bedaquiline, CFZ = clofazimine, DLM = delamanid, LZD = linezolid, ETH = ethionamide, RFB = rifabutin.

Of the 12,289 isolates with matched phenotypic and genotypic data, 6,814 (55.4%) were resistant to at least one drug (Fig. 3b). For the purpose of describing five broader resistance categories present in the dataset, we assumed that all MICs that could not be read had susceptible phenotypes. These five resistance categories comprise: isoniazid and rifampicin susceptible with resistance to another antitubercular drug, isoniazid resistant but rifampicin susceptible, RR/MDR, pre-XDR (RR/MDR + fluoroquinolone resistance), and XDR (RR/MDR + fluoroquinolone resistance + resistance to a group A agent: bedaquiline or linezolid)). Consequently, the calculated prevalence of MDR, XDR *etc*. in the dataset (Fig. 3) are likely under- estimates. Of the isolates resistant to one or more drugs, 22.8% were resistant to isoniazid and not rifampicin, 68.8% were either RR or MDR and 8.4% were resistant to at least one antitubercular drug, but *not* isoniazid or rifampicin (Fig. 3b). Of the RR/MDR isolates, 38.8% were pre-XDR and 3.0% were XDR. Two of the XDR isolates returned a resistant phenotype to all 13 of the drugs assayed (Table S5) and therefore could be reasonably described as totally drug resistant (TDR). One such isolate belonged to L4 and was contributed by South Africa, and the other belonged to L2 with an unknown country of origin contributed by Sweden.

The proportion of drug susceptible isolates collected differed between countries, as not all laboratories oversampled for resistance (Fig. 3c). In countries that contributed more than 100 resistant isolates, each of the broad phenotypic resistance categories in Fig. 3b were seen except for Peru, Vietnam and Nepal which did not contribute any XDR isolates (Fig. 3c). Vietnam and Brazil sampled a high proportion of non-MDR/RR resistant phenotypes; 73.9% and 55.1% of resistant isolates contributed by these countries, respectively, were neither MDR nor RR. For Nepal and India, an especially high proportion of the MDR/RR isolates contributed were fluoroquinolone resistant (92.9% and 69.8% respectively), which has been previously observed for this geographical region (32, 33).

Of the 6814 resistant isolates, 256 were from L1, 2886 were from L2, 625 were from L3 and 3043 were from L4. All five broader categories of resistance were represented in the four major *M. tuberculosis* lineages (Fig. 3d). We note that the relative proportions of resistance categories will have been influenced by the different local sampling approaches since lineage distributions are typically geographically distinct (Fig. S2). Bearing this in mind, we observe that in the compendium, L3 isolates contained the most MDR/RR isolates as a proportion of resistant isolates (77.6%), L2 isolates contained the most pre-XDR isolates as a proportion of MDR/RR isolates (54.2%) and L2 contained the most XDR isolates as a proportion of MDR/RR isolates (4.7%).

### Co-occurrence of drug resistance amongst the CRyPTIC isolates

As we measured MICs to 13 drugs in parallel, we can ask whether, and how often, co-occurrence of drug resistance occurs amongst the isolates. We found isolates with all possible two-drug resistant combinations in this dataset (Fig. 4a, Table S6). With the exception of correlations between drugs in the same class (rifabutin vs rifampicin, moxifloxacin vs levofloxacin), Isoniazid resistance was the most strongly associated with resistance to each of the other drugs. Resistance to any of the drugs was also strongly associated with resistance to rifampicin. Of the second line drugs, levofloxacin and moxifloxacin were more commonly seen as a second resistant phenotype than the injectable drugs kanamycin and amikacin.

**Figure 4:**
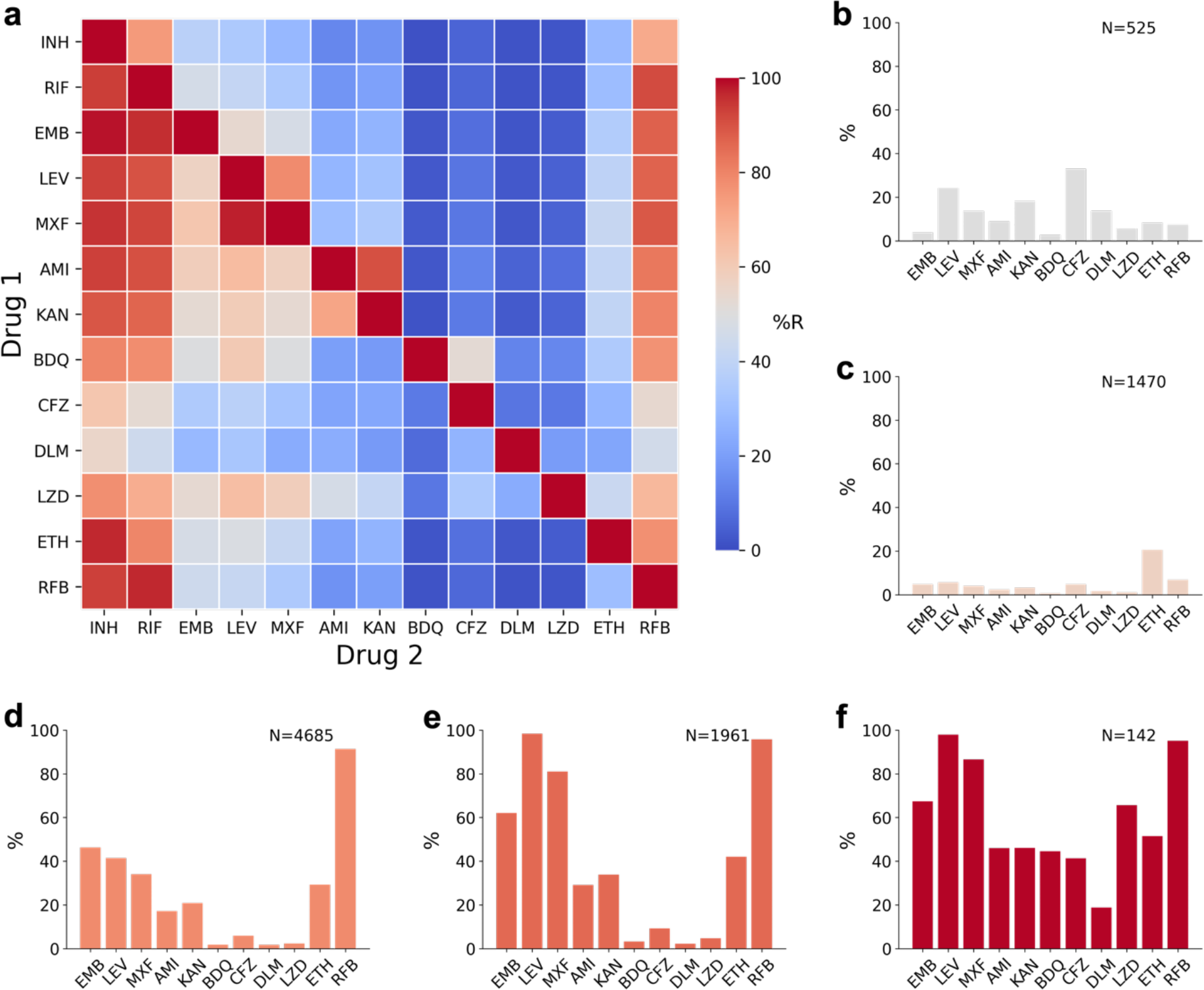
Co-occurrence of resistance to one drug conditional on resistance to another drug, or to resistance background. **a)** The heatmap shows the probability of an isolate being resistant to Drug 2 if it is resistant to Drug 1, percentages are given in Table S4. **(b-g)** Percentage of isolates that are resistant to another of the 13 drugs in a background of **(b)** isoniazid susceptible + rifampicin susceptible (but resistant to at least one other antitubercular drug), **(c)** isoniazid resistant + rifampicin susceptible, **(d)** MDR/RR **(e)** Pre-XDR, **(f)** XDR. Only samples with definite phenotypes for RIF in MDR backgrounds and RIF and INH in non-MDR backgrounds and the additional drug are included. Drug acronyms: INH = isoniazid, RIF = rifampicin, EMB = ethambutol, LEV = levofloxacin, MXF = moxifloxacin, AMI = amikacin, KAN = kanamycin, BDQ = bedaquiline, CFZ = clofazimine, DLM = delamanid, LZD = linezolid, ETH = ethionamide, RFB = rifabutin.

Resistance to both drugs in the aminoglycoside class was common in the dataset; 90.4% of amikacin resistant isolates were also resistant to kanamycin although significantly fewer kanamycin resistant isolates were resistant to amikacin (72.0%, *p* <0.00001) (Fig. 4a). In a similar fashion, a smaller proportion of rifampicin resistant isolates were resistant to rifabutin than rifabutin resistant isolates that were resistant to rifampicin (91.3%, 96.8%, *p* <0.00001) while a smaller proportion of levofloxacin resistant isolates were resistant to moxifloxacin than moxifloxacin resistant isolates that were resistant to levofloxacin (78.5%, 97.6%, *p* <0.00001).

Differences in drugs of the same class are also well documented by *in vitro* studies (34–36).

Isolates resistant to the NRDs bedaquiline, clofazimine, delamanid and linezolid were most likely to also be resistant to isoniazid, followed by rifampicin and rifabutin. The NRDs were less commonly seen as a second resistance phenotype and the smallest proportional resistance combinations involved the NRDs (*e.g.* 1.5% of isoniazid resistant isolates were bedaquiline resistant). Within the NRDs however, co-occurrence of resistance was proportionally higher; bedaquiline, linezolid and delamanid resistance was commonly seen with clofazimine resistance (52.4%, 34.2% and 26.3% of isolates having co-resistance with clofazimine respectively).

### Additional antibiotic resistance in isolates with non-MDR or MDR phenotypic backgrounds

To further investigate drug resistance patterns amongst the isolates, we examined in more detail the correlation structure of phenotypes by conditioning on different resistance backgrounds including isoniazid and rifampicin susceptible, isoniazid resistant and rifampicin susceptible, rifampicin resistant and isoniazid susceptible, MDR, pre-XDR and XDR (Fig. 4b-f). We found that a greater proportion of isolates that were susceptible to isoniazid and rifampicin were resistant to the second line drugs levofloxacin (24.1%), kanamycin (18.1%), moxifloxacin (13.7%), and amikacin (8.9%) than the first line drug ethambutol (3.8%) (Fig. 4b). The proportion of isolates resistant to clofazimine or levofloxacin was particularly high (32.9% and 24.1%, respectively), and more isolates were resistant to these two drugs than ethambutol in an isoniazid resistant and rifampicin susceptible background but not in MDR/RR isolates (Fig. 4c-f).

MDR/RR isolates were most commonly resistant (excluding rifabutin) to the first line drug ethambutol (46.3%), closely followed by levofloxacin (41.4%). As expected, the proportion of fluoroquinolone resistance was higher in MDR/RR isolates than non-MDR isolates (37) and we found a greater proportion of isolates were resistant to levofloxacin than moxifloxacin, a pattern seen in all other backgrounds (Fig. 4c-f). For the aminoglycosides, a greater percentage of MDR/RR isolates were kanamycin resistant (21.8%) than amikacin resistant (18.1%), a trend seen in all other backgrounds.

For isolates with an XDR phenotype, a higher proportion were resistant to linezolid than bedaquiline (66.7% compared to 44.6%) and 11.3% of XDR isolates were resistant to both bedaquiline and linezolid (Fig. 4f). XDR isolates were also resistant to the other NRDs, clofazimine (41.3%) and delamanid (18.8%). In non-XDR backgrounds the most common NRD resistance seen was clofazimine (Fig. 4b-e).

### Genetic-based predictions of resistance

To establish a baseline measure of how well resistance and susceptibility could be predicted based on the state of the art prior to the CRyPTIC project, we compared genetic-based predictions of susceptibility and resistance to the binary phenotypes derived from MICs for eight drugs and for all isolates in this compendium (Table 2). Since these data were not collected prospectively or randomly, but indeed are enriched for resistance, the calculated error rates are not representative of how well such a method would perform in routine clinical use. The results were broadly in line with prior measurements on a smaller (independent) set (23). The hybrid catalogue does not make predictions for rifabutin, linezolid, bedaquiline, delamanid or clofazimine; indeed, this is one of the main aims of the consortium and new catalogues published by CRyPTIC and the WHO will begin to address this shortcoming (Fig. 5) (26).

**Figure 5:**
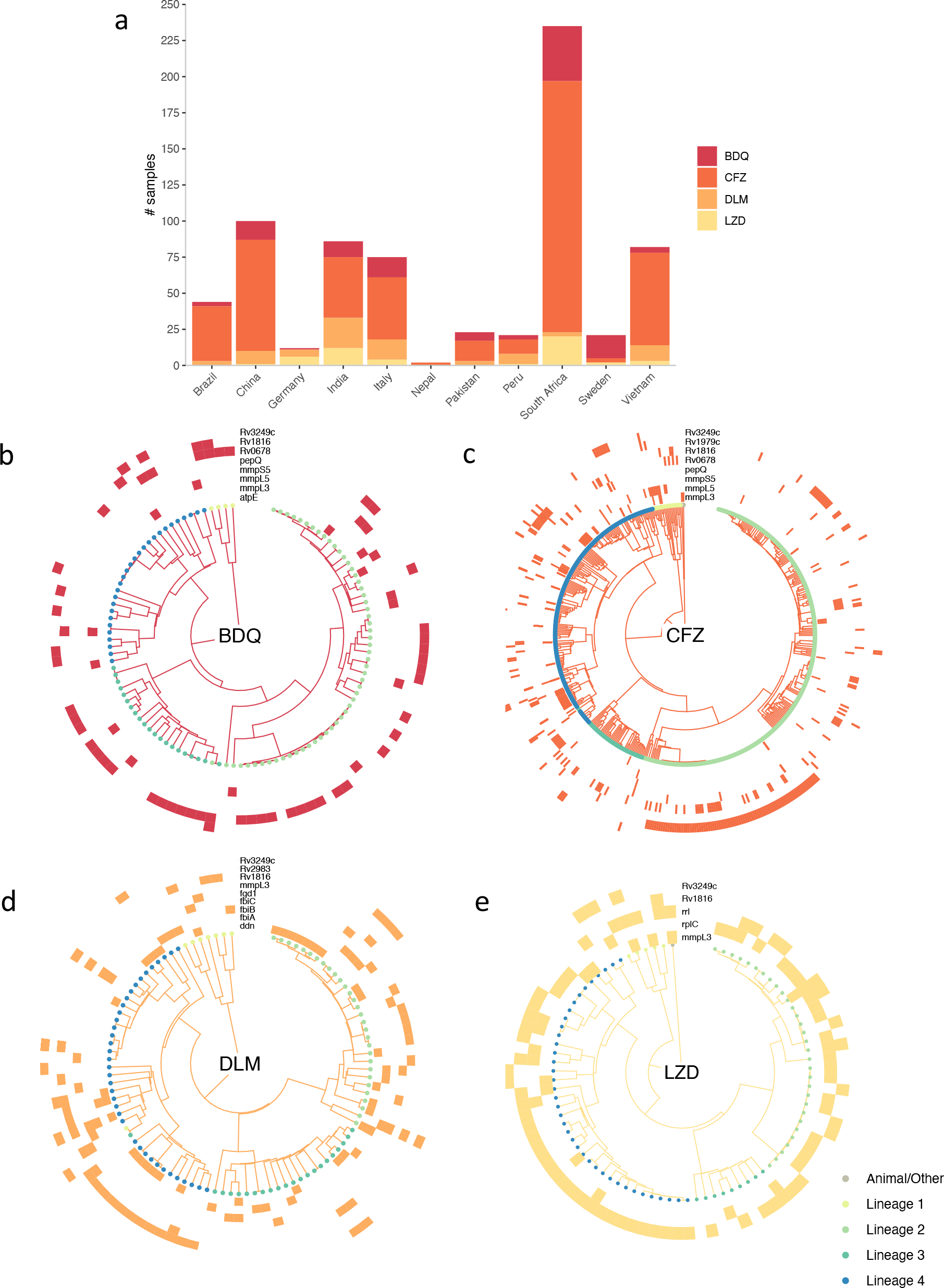
Resistance to bedaquiline, clofazimine, delamanid and linezolid amongst M. tuberculosis CRyPTIC isolates. **a)** The prevalence (within these data) of resistance to bedaquiline (BDQ), clofazimine (CFZ), delamanid (DLM) and linezolid (LZD) per country or origin or collection site. Phylotrees are shown for isolates phenotypically resistant to b) BDQ, c) CFZ, d) DLM and e) LZD. Tip point colours denote lineage. Each outer track represents a gene thought to be associated with resistance and coloured blocks denote the presence of a non-synonymous mutation in the relevant gene for a given isolate. Mutations in these genes that are either associated with sensitivity or present in >5% of the collection of isolates as a whole were ignored.

**Table 2:** Predicting phenotypic resistance using genetics Statistics on how well phenotypic resistance could be predicted using a standard resistance catalogue that predates the CRyPTIC project. TP: the number of phenotypically resistant samples are that correctly identified as resistant (“true positives”); FP, the number of phenotypically susceptible samples that are falsely identified as resistant (“false positives”); TN, the number of phenotypically susceptible samples that are correctly identified as susceptible (“true negatives”); FN, the number of phenotypically resistant samples that are incorrectly identified as susceptible (“false negative”); VME, very major error rate (false-negative rate), 0-1; ME, major error rate (false-positive rate), 0-1; PPV, positive predictive value, 0-1; NPV, negative predictive value, 0-1. Drug acronyms: INH = isoniazid, RIF = rifampicin, EMB = ethambutol, AMI = amikacin, KAN = kanamycin, LEV = levofloxacin, MXF = moxifloxacin, ETH = ethionamide.

|  | TP | FP | TN | FN | VME | ME | PPV | NPV |
| --- | --- | --- | --- | --- | --- | --- | --- | --- |
| **INH** | 5493 | 142 | 5622 | 224 | 0.039 | 0.025 | 0.961 | 0.975 |
| **RIF** | 4535 | 435 | 6669 | 107 | 0.023 | 0.061 | 0.977 | 0.939 |
| **EMB** | 1919 | 513 | 6702 | 111 | 0.055 | 0.071 | 0.945 | 0.929 |
| **LEV** | 1689 | 255 | 8104 | 184 | 0.098 | 0.031 | 0.902 | 0.969 |
| **MXF** | 1358 | 504 | 9022 | 160 | 0.105 | 0.053 | 0.895 | 0.947 |
| **AMI** | 632 | 84 | 10117 | 163 | 0.205 | 0.008 | 0.795 | 0.992 |
| **KAN** | 735 | 124 | 9043 | 197 | 0.211 | 0.014 | 0.789 | 0.986 |
| **ETH** | 971 | 114 | 9183 | 511 | 0.345 | 0.012 | 0.655 | 0.988 |

Table 3 shows the top mutations found amongst isolates phenotypically resistant to first- and second- line drugs. As expected, *rpoB* S450L was the most prevalent mutation associated with rifampicin resistance and *katG* S315T was the most prevalent mutation associated with isoniazid resistance. Mutations in *gyrA* dominate amongst fluoroquinolone resistant isolates; D94G and A90V are the two most frequently occurring mutations for levofloxacin and moxifloxacin.

**Table 3:** The top mutations associated with phenotypic drug resistance. Depicted is a survey of the resistance-associated mutations present in CRyPTIC isolates (7, 25). “VARIANT”: non-synonymous amino acid mutations are denoted by upper case letters while nucleotide substitutions for non-coding sequences are denoted by lower case letters. Negative numbers denote substitutions in promoter regions; “GENE”: genic region of interest in which “Variant” can be found; “*n”*: number of phenotypically resistant isolates with ”VARIANT”; “%”: percentage of total phenotypically resistant isolates with “VARIANT”.

|  | GENE | VARIANT | *n* | % |
| --- | --- | --- | --- | --- |
| **Rifampicin** | *rpoB* | S450L | 2914 | 62.2 |
|  | *rpoB* | D435V | 506 | 10.8 |
|  | *rpoB* | H445D | 202 | 4.3 |
|  | *rpoB* | H445Y | 147 | 3.1 |
|  | *rpoB* | D435Y | 112 | 2.4 |
| **Isoniazid** | *katG* | S315T | 3668 | 70.0 |
|  | *fabG1* | c-15t | 829 | 15.8 |
|  | *fabG1* | g-17t | 176 | 3.4 |
|  | *fabG1* | t-8c | 154 | 2.9 |
|  | *inhA* | I194T | 56 | 1.1 |
| **Ethambutol** | *embB* | M306V | 1131 | 50.0 |
|  | *embB* | M306I | 1001 | 44.3 |
|  | *embB* | Q497R | 449 | 19.9 |
|  | *embB* | G406A | 164 | 7.2 |
|  | *embB* | G406D | 105 | 4.6 |
| **Kanamycin** | *rrs* | a1401g | 660 | 58.9 |
|  | *eis* | c-14t | 70 | 6.2 |
|  | *eis* | g-10a | 53 | 4.7 |
| **Amikacin** | *rrs* | a1401g | 660 | 74.7 |
|  | *rrs* | g1484t | 7 | 0.8 |
| **Levofloxacin** | *gyrA* | D94G | 783 | 36.5 |
|  | *gyrA* | A90V | 487 | 22.7 |
|  | *gyrA* | D94N | 157 | 7.3 |
|  | *gyrA* | D94A | 133 | 6.2 |
|  | *gyrA* | S91P | 92 | 4.3 |
| **Moxifloxacin** | *gyrA* | D94G | 783 | 45.4 |
|  | *gyrA* | A90V | 487 | 28.2 |
|  | *gyrA* | D94N | 157 | 9.1 |
|  | *gyrA* | D94A | 133 | 7.7 |
|  | *gyrA* | D94Y | 70 | 4.1 |
| **Ethionamide** | *fabG1* | c-15t | 829 | 48.0 |
|  | *fabG1* | L203L | 124 | 7.2 |

### Resistance to new and re-purposed drugs

As previously stated, relatively few isolates are resistant to the NRDs, bedaquiline (*n* = 109), clofazimine (*n* = 525), delamanid (*n* = 186) and linezolid (*n* = 156). South Africa contributed the greatest number of isolates resistant to bedaquiline, clofazimine and linezolid (Fig. 5a), while China and India contributed the most isolates resistant to delamanid. Since the collection protocol differed between laboratories it is not possible to infer any differences in the relative prevalence of resistance to the NRDs in these countries. The results of a survey of all non-synonymous mutations in genes known or suspected to be involved in resistance to these four drugs (*e.g. rv0678, mmpL5, pepQ, ddn, rplC, rrl etc.*) are depicted in Fig. 5b-e (38–42). Mutations known to be associated with sensitivity were ignored, along with mutations that occurred at a frequency of ≥ 5% amongst all isolates (as 0.05% of total isolates are resistant to the NRDs). In contrast to first- and second-line drugs, there are no mutations within a single gene/small group of genes that can fully explain resistance to any NRD, and indeed no single gene (if there were, they would be visible as complete rings around the phylotrees in figure 5). Note that the role of most of these mutations in resistance remains undetermined.

### Case study on rifampicin mono-resistance

Around 1% of TB cases are rifampicin mono-resistant (RMR) and the frequency is increasing (1, 43). The WHO does not recommend isoniazid for RMR treatment, despite it being effective; this is likely due to the reliance on assays such as Xpert® MTB/RIF which cannot distinguish between RMR and MDR. Use of isoniazid could improve treatment outcomes for RMR patients which are currently similar to that of MDR TB, including a higher risk of death compared to drug susceptible infections (44, 45). Due to its low natural prevalence, RMR has been poorly studied to date but increasingly large clinical TB datasets, such as the one presented here, make its study now feasible.

For this case study, we defined RMR as any isolate that is rifampicin resistant and isoniazid susceptible, and discounted isolates with no definite phenotype for either drug. Of the 4,655 rifampicin resistant isolates in the compendium that also had a phenotype for isoniazid, 302 (6.5%) were RMR. These isolates were contributed by 12 different countries, and we found South African and Nigerian contributions had a significantly higher proportion of RMR isolates than that of the total dataset at 17.5% (*p* <0.00001) and 27.3% (*p* = 0.00534) respectively (Fig. 6a) compared with 6.5% for the total dataset. We note that these proportions are influenced by sampling strategies but the higher contribution of RMR isolates from South Africa is consistent with previous studies (43).

**Figure 6:**
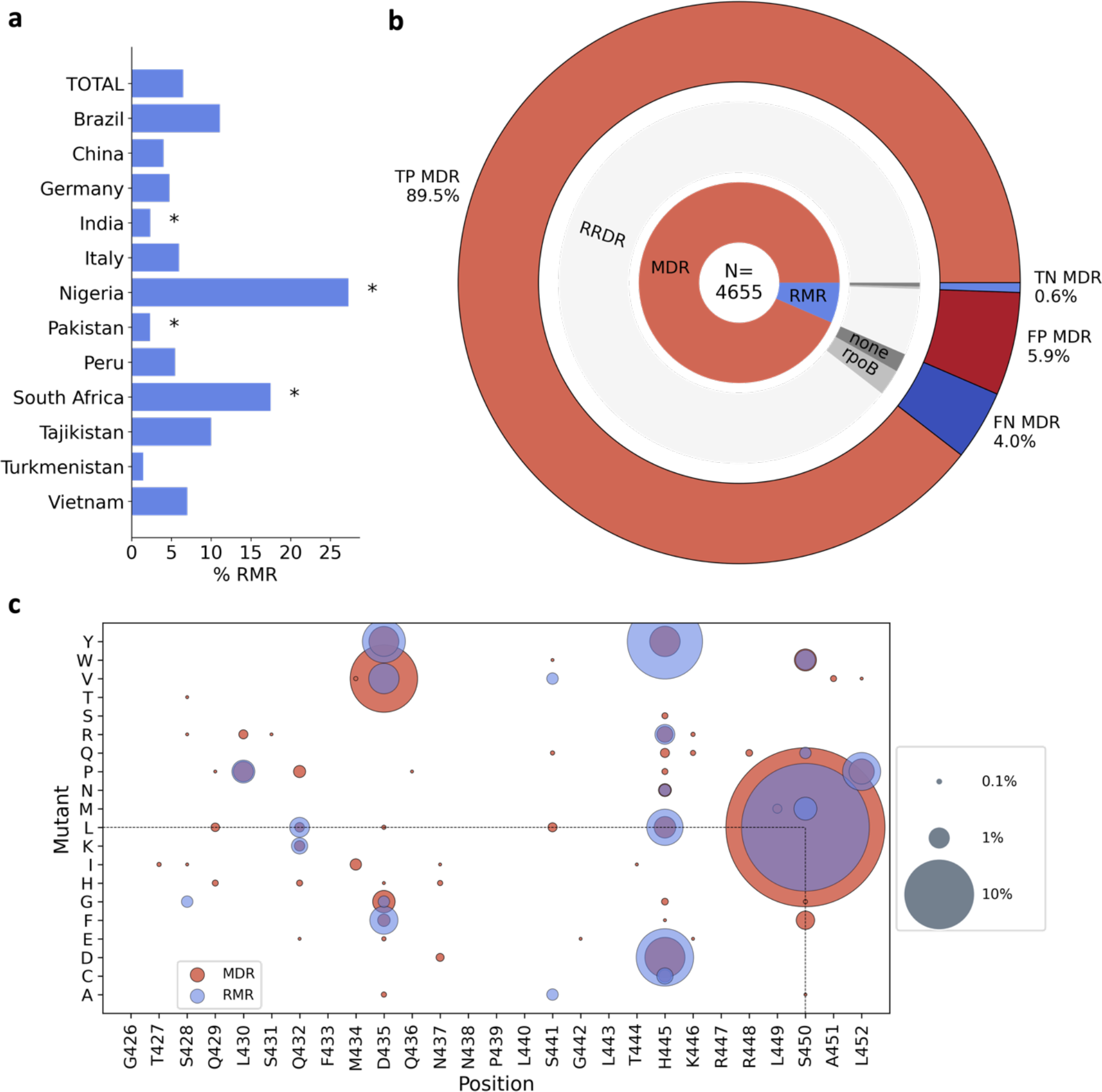
Rifampicin mono resistance. **a)** Percentage of rifampicin resistant isolates that are rifampicin mono-resistant (RMR) by country of isolate origin. * indicates RMR proportions that were significantly different from that of the total dataset using a two tailed z-test with 95% confidence. **b)** MDR predictions for rifampicin resistant isolates made using the Xpert® MTB/RIF assay proxy. N is the total number of rifampicin resistant isolates. The inner ring shows the proportion of rifampicin resistant isolates that are MDR and RMR. The middle ring represents the proportions of MDR and RMR isolates that have a SNP (synonymous or non-synonymous) in the RRDR of *rpoB* (RRDR), no RRDR SNP but a SNP elsewhere in the *rpoB* gene (rpoB) and no *rpoB* mutations (none). The outer ring shows the expected true positive (TP), true negative (TN), false positive (FP) and false negative (FN) MDR predictions of Xpert® MTB/RIF assay, based on the SNPs present in the rifampicin resistant isolates. **c)** Non-synonymous mutations found in the RRDR of *rpoB* in RMR isolates and MDR isolates. Presence of a coloured spot indicates that the mutation was found in RMR/MDR isolates and spot size corresponds to the proportion of RMR or MDR isolates carrying that mutation.

### Rifampicin mono-resistance is incorrectly predicted by current diagnostics

A widely used, WHO-endorsed diagnostic tool, the Xpert® MTB/RIF assay, uses a proxy whereby *any* SNP detected in the “rifampicin-resistance determining region” (RRDR) of *rpoB* results in a prediction of MDR. However, the suitability of the proxy is dependent upon prevalence of RMR in the population (43). We tested the reliability of this on the 4,655 rifampicin resistant isolates in our dataset that had a phenotype for isoniazid (Fig. 6b).

Of these isolates, 4,353 (93.5%) were MDR and 302 (6.5%) were RMR. 187 of the MDR isolates had no RRDR mutation and therefore 4.0% of isolates in this study would be predicted as false negative MDR by the Xpert® MTB/RIF assay. 276 of the RMR isolates had a mutation in the RRDR of *rpoB* and so the Xpert® MTB/RIF assay proxy would incorrectly predict 5.9% of the rifampicin resistant isolates as false positive MDR cases. However, overall, the Xpert® MTB/RIF assay proxy correctly predicts 89.5% of the rifampicin resistant isolates as MDR and 0.6% of the isolates as non-MDR in this dataset, which suggests it is a reasonably successful diagnostic tool with > 90% accuracy for MDR classification of rifampicin resistant isolates. As our dataset is oversampled for resistance, it likely contains a higher prevalence of RMR than the global average and hence the Xpert® MTB/RIF assay is likely to perform better on more representative data. However, the analysis shows how the increasing global levels of RMR TB cases could increase the number of false positive MDR diagnoses by the Xpert® MTB/RIF assay, denying isoniazid treatment to a greater number of patients who would then be moved on to less effective drugs.

### There are genetic differences between rifampicin mono-resistant and multidrug resistant isolates

We have analysed our matched phenotypic and genotypic data to examine whether there were any differences in the genetic determinants of rifampicin resistance between RMR and MDR isolates as was seen in a recent study of South African isolates (46). The proportion of RMR isolates with no *rpoB* mutation (5.3%, Fig. 6b) was significantly higher than that of MDR isolates (1.8%, *p* <0.00001). This suggests that non-target-mediated resistance mechanisms, such as upregulation of rifampicin specific efflux pumps, could be more influential in providing protection against rifampicin in RMR isolates than in MDR isolates.

The majority of RMR and MDR isolates contained one or more SNPs in *rpoB*, with the most having at least one mutation in the RRDR. To date, several non-synonymous RRDR mutations have been found in RMR *M. tuberculosis* isolates, including the resistance conferring mutations S450L, H445D and D435Y, which are also seen in MDR isolates (47, 48). For both RMR and MDR isolates in this dataset, the most common *rpoB* RRDR mutation seen was S450L (63.6% and 41.1% of isolates respectively, Fig. 6c). Five mutations were present in RMR isolates that were not seen in MDR isolates: S428G, S441A, S441V, S450M and S450Q, however these were seen at low prevalence (< 2%) of RMR isolates. We found more RMR isolates had His445 mutated than MDR isolates (27.8% of RMR and 9.5% of MDR, *p* <0.00001), and mutations at Ser450 and Asp435 were more prevalent in MDR isolates than RMR isolates (43.7% of RMR and 65.8% of MDR (*p* <0.00001), and 9.3% of RMR and 15.5% of MDR (*p* = 0.00328) respectively).

In RMR isolates we observed 27 different *rpoB* mutations that fall outside the RRDR; 11 were found in RMR but not MDR isolates and all were seen at < 2% prevalence (Fig. S6). The most common non-RRDR mutation in both MDR and RMR isolates was a cytosine to thymine mutation 61 bases upstream of the *rpoB* start codon (10.1% and 8.6% of isolates respectively). The resistance conferring mutations, *rpoB* I491F, V695L and V170F, were seen at low proportions (< 2% of isolates) with no significant difference between MDR and RMR isolates.

## Discussion

This compendium of *M. tuberculosis* clinical isolates is the result of an extensive global effort by the CRyPTIC consortium to better map the genetic variation associated with drug resistance. Through its sheer size and by oversampling for resistance, the compendium gives an unparalleled view of resistance and resistance patterns amongst the panel of 13 antitubercular compounds studied. This study serves to summarise the data within the compendium and to highlight the existence of the open-access resource to the wider community to help better inform future treatment guidelines and steer the development of improved diagnostics.

Starting with first-line drugs, molecular based diagnostic assays have vastly improved the detection of and the speed at which we find drug resistant TB cases, resulting in improved quality of care for patients. However, relying solely on these diagnostic methods has several drawbacks. Aside from the Xpert® MTB/RIF assay potentially increasing false positive MDR diagnoses as discussed earlier in the RMR case study, the assay assumes isoniazid resistance upon detection of rifampicin resistance. Thus, less is known about the prevalence of mono isoniazid resistance or ’true’ cases of MDR (confirmed rifampicin *and* isoniazid resistance) (1) and with large datasets such as this compendium, we can further investigate these important and rarer clinical phenotypes (like that of RMR in our case study). Another example of a rarer phenotype is that of isoniazid-resistant and rifampicin-susceptible (Hr-TB) isolates; a greater number of these were contributed by CRyPTIC countries than RMR isolates *(n* = 1470 versus *n* = 302), a pattern also recently observed in a global prevalence study (49). A modified 6-month treatment regimen is now recommended for Hr-TB (rifampicin, ethambutol, levofloxacin and pyrazinamide), and as a result of inadequate diagnosis many of the 1.4 million global Hr-TB estimated cases would have received inadequate and unnecessarily longer treatment regimens (1, 42). Encouragingly, CRyPTIC isolates with a Hr-TB background exhibited relatively low levels of resistance to other antitubercular drugs, including those in the augmented regimen (Fig. 4c). However, without appropriate tools to assess and survey this, we will continue to misdiagnose and infectively treat these clinical cases. In 2018, CRyPTIC and the 100,000 Genomes project demonstrated that genotypic prediction from WGS correlates well with culture-based phenotype for first-line drugs, which is reflected in our summary of the genetic catalogue applied to this dataset (Table 3) (7). While predictions can be made to a high level of sensitivity and specificity, there is still more to learn, as exemplified by the isolates in the compendium that despite being resistant to rifampicin and isoniazid could not be described genetically (Table 2). This shortfall, along with the limitations of molecular based diagnostic assays, highlights the need for continual genetic surveillance and shines a favourable light on a WGS- led approach.

A strength of this compendium lies with the data collated for second-line drugs. A greater proportion of drug resistant isolates had additional resistance to fluoroquinolones than second line injectable drugs (Fig. 4a). This could be due to more widespread use of fluoroquinolones as well as their ease of administration and hence them being recommended over injectables for longer MDR treatment regimens (1). Concerningly, we found that resistance to levofloxacin and moxifloxacin, and kanamycin and amikacin, were more common than resistance to the mycobacterial specific drug ethambutol in an isoniazid and rifampicin susceptible background (Fig. 4b), suggesting a level of pre-existing resistance to second-line drugs. This concurs with a systematic review that found patients previously prescribed fluoroquinolones were three times more likely to have fluoroquinolone resistant TB (50). Careful stewardship of fluoroquinolones, both in TB and other infectious diseases, will be paramount for the success of treatment regimens. Despite variability in sample collection, we observed high proportions of fluoroquinolone resistant MDR/RR isolates from some countries and therefore suggest that MDR treatment regimens could be improved by optimisation on a geographic basis.

Further treatment improvement could also be made by the selection of appropriate drugs from each class. For example, the WHO recommends switching from kanamycin to amikacin when treating MDR TB patients (51) and the compendium supports this recommendation as we saw more resistance to kanamycin than amikacin in all phenotypic backgrounds. For fluoroquinolones, more isolates were resistant to levofloxacin than moxifloxacin in all phenotypic backgrounds suggesting moxifloxacin may by the most appropriate fluoroquinolone to recommend, although we note this conclusion is critically dependent on the validity of the cutoff, here an ECOFF, used to infer resistance. However, the amenability of drugs to catalogue-based genetic diagnostics is also an important consideration, and our data suggest levofloxacin resistance could be predicted more reliably than moxifloxacin, with fewer false positives predicted (Table 2). Testing for fluoroquinolone resistance using molecular diagnostic tests remains limited. Global data from the past 15 years suggests that the proportion of MDR/RR TB cases resistant to fluoroquinolones sits at around 20%, with these cases primarily found in regions of high MDR-TB burden (1). While recently approved tools, such as the Cepheid Xpert® MTB/XDR cartridge, will permit both isoniazid and fluoroquinolone testing to be increased, the same pitfalls are to be encountered regarding targeted diagnostic assays (52). In contrast, the genetic survey in this study demonstrates the potential of WGS for genetic prediction of resistance to second-line drugs and studies within the consortium to investigate this are underway.

The CRyPTIC compendium has facilitated the first global survey of resistance to NRDs. Reassuringly, prevalence of resistance to the NRDs was substantially lower than for first- and second-line agents in the dataset (Fig. 3a), and resistance to the new drugs bedaquiline and delamanid was less common than the repurposed drugs clofazimine and linezolid in an MDR/RR background (Fig. 4c). However, the presence of higher levels of delamanid and clofazimine resistance than ethambutol resistance in the isoniazid and rifampicin susceptible background does suggest some pre-existing propensity towards NRD resistance (Fig. 4b).

Co-resistance between NRDs was seen in isolates in the compendium, the most common being isolates resistant to both bedaquiline and clofazimine. This link is well documented and has been attributed to shared resistance mechanisms such as non-synonymous mutations in *Rv0678* which were found in both clofazimine and bedaquiline resistant isolates in the compendium (42) (Fig. 5b,c). Increased clofazimine use could further increase the prevalence of *M. tuberculosis* isolates with clofazimine and bedaquiline co-resistance, limiting MDR treatment options including using bedaquiline as the backbone of a shorter MDR regimen (53). Therefore, proposed usage of clofazimine for other infectious diseases should be carefully considered.

The WHO recommends against the use of bedaquiline and delamanid in combination to prevent the development of co-resistance, which could occur relatively quickly (54); the rate of spontaneous evolution of delamanid resistance *in vitro* has been shown to be comparable to that of isoniazid, and likewise bedaquiline resistance arises at a comparable rate to rifampicin resistance (55). In this compendium, 12.9% of bedaquiline resistant isolates were resistant to delamanid and 7.1% of delamanid resistant isolates were resistant to bedaquiline. Several scenarios could account for this, including the presence of shared resistance mechanisms. For example, as bedaquiline targets energy metabolism within the cell, changes to cope with energy/nutrient imbalances upon the acquisition of resistance-associated ATPase pump mutations may result in cross resistance to delamanid in a yet unknown or unexplored mechanism (12). it is imperative that genetic determinants of resistance are fully explored for the NRDs, as these are our current treatments of last resort, with special attention given to those mechanisms that could be shared with other agents.

In the meantime, careful stewardship and phenotypic and genotypic surveillance of the NRDs should be implemented, including linezolid and clofazimine which are now group A and B drugs respectively for MDR treatment (1).

Several research avenues are being actively explored by the CRyPTIC consortium that make further use of this compendium, including: *i)* relating genetic mutations to quantitative changes in the minimum inhibitory concentrations of different drugs (12); *ii)* genome-wide association studies (14); *iii)* training machine learning models that can predict resistance (13); *iv*) exploration of the genetic determinants of resistance to second line and NRDs (15). Collectively these studies share the same aim of facilitating the implementation of WGS-directed resistance prediction in the clinic. Finally, we urge other researchers to explore and analyse this large dataset of *M. tuberculosis* clinical isolates and hope it will lead to a wave of new and inciteful studies that will positively serve the TB community for years to come.

## CRyPTIC Consortium Members

Alice Brankin5,*,** and Kerri M Malone23,*,**, Ivan Barilar1, Simone Battaglia2, Emanuele Borroni2, Angela Pires Brandao3,4, Andrea Maurizio Cabibbe2, Joshua Carter6, Darren Chetty7, Daniela Maria Cirillo2, Pauline Claxton8, David A Clifton5, Ted Cohen9, Jorge Coronel10, Derrick W Crook5, Viola Dreyer1, Sarah G Earle5, Vincent Escuyer11, Lucilaine Ferrazoli4, George Fu Gao12, Jennifer Gardy13, Saheer Gharbia14, Kelen Teixeira Ghisi4, Arash Ghodousi2,15, Ana Lúıza Gibertoni Cruz5, Louis Grandjean16, Clara Grazian17, Ramona Groenheit18, Jennifer L Guthrie19,20, Wencong He12, Harald Hoffmann21,22, Sarah J Hoosdally5, Martin Hunt23,5, Nazir Ahmed Ismail24, Lisa Jarrett25, Lavania Joseph24, Ruwen Jou26, Priti Kambli27, Rukhsar Khot27, Jeff Knaggs23,5, Anastasia Koch28, Donna Kohlerschmidt11, Samaneh Kouchaki5,29, Alexander S Lachapelle5, Ajit Lalvani30, Simon Grandjean Lapierre31, Ian F Laurenson8, Brice Letcher23, Wan-Hsuan Lin26, Chunfa Liu12, Dongxin Liu12, Ayan Mandal32, Mikael Mansjo18, Daniela Matias25, Graeme Meintjes28, Flávia de Freitas Mendes4, Matthias Merker1, Marina Mihalic22, James Millard7, Paolo Miotto2, Nerges Mistry32, David Moore33,10, Kimberlee A Musser11, Dumisani Ngcamu24, Hoang Ngoc Nhung34, Stefan Niemann1,35, Kayzad Soli Nilgiriwala32, Camus Nimmo16, Max O’Donnell36, Nana Okozi24, Rosangela Siqueira Oliveira4, Shaheed Vally Omar24, Nicholas Paton37, Timothy EA Peto5, Juliana Maira Watanabe Pinhata4, Sara Plesnik22, Zully M Puyen38, Marie Sylvianne Rabodoarivelo39, Niaina Rakotosamimanana39, Paola MV Rancoita15, Priti Rathod25, Esther Robinson25, Gillian Rodger5, Camilla Rodrigues27, Timothy C Rodwell40,41, Aysha Roohi5, David Santos-Lazaro38, Sanchi Shah32, Thomas Andreas Kohl1, Grace Smith25,14, Walter Solano10, Andrea Spitaleri2,15, Philip Supply42, Adrie JC Steyn7, Utkarsha Surve27, Sabira Tahseen43, Nguyen Thuy Thuong Thuong34, Guy Thwaites34,5, Katharina Todt22, Alberto Trovato2, Christian Utpatel1, Annelies Van Rie44, Srinivasan Vijay45, Timothy M Walker5,34, A Sarah Walker5, Robin Warren46, Jim Werngren18, Maria Wijkander18, Robert J Wilkinson47,48,30, Daniel J Wilson5, Penelope Wintringer23, Yu-Xin Xiao26, Yang Yang5, Zhao Yanlin12, Shen-Yuan Yao24, Baoli Zhu49, Philip W Fowler5, Zamin Iqbal **

*equal contribution authors

**co-corresponding authors

### Institutions

1Research Center Borstel, Borstel, Germany

2IRCCS San Raffaele Scientific Institute, Milan, Italy

3Oswaldo Cruz Foundation, Rio de Janeiro, Brazil

4Institute Adolfo Lutz, Sào Paulo, Brazil

5University of Oxford, Oxford, UK

6Stanford University School of Medicine, Stanford, USA

7Africa Health Research Institute, Durban, South Africa

8Scottish Mycobacteria Reference Laboratory, Edinburgh, UK

9Yale School of Public Health, Yale, USA

10Universidad Peruana Cayetano Heredia, Lima, Peru

11Wadsworth Center, New York State Department of Health, Albany, USA

12Chinese Center for Disease Control and Prevention, Beijing, China

13Bill & Melinda Gates Foundation, Seattle, USA

14UK Health Security Agency, London, UK

15Vita-Salute San Raffaele University, Milan, Italy

16University College London, London, UK

17University of New South Wales, Sydney, Australia

18Public Health Agency of Sweden, Solna, Sweden

19The University of British Columbia, Vancouver, Canada

20Public Health Ontario, Toronto, Canada

21SYNLAB Gauting, Munich, Germany

22Institute of Microbiology and Laboratory Medicine, IMLred, WHO-SRL Gauting, Germany

23EMBL-EBI, Hinxton, UK

24National Institute for Communicable Diseases, Johannesburg, South Africa

25UK Health Security Agency, Birmingham, UK

26Taiwan Centers for Disease Control, Taipei, Taiwan

27Hinduja Hospital, Mumbai, India

28University of Cape Town, Cape Town, South Africa

29University of Surrey, Guildford, UK

30Imperial College, London, UK

31Université de Montréal, Canada

32The Foundation for Medical Research, Mumbai, India

33London School of Hygiene and Tropical Medicine, London, UK

34Oxford University Clinical Research Unit, Ho Chi Minh City, Viet Nam

35German Center for Infection Research (DZIF), Hamburg-Lübeck-Borstel-Riems, Germany

36Colombia University Irving Medical Center, New York, USA

37National University of Singapore, Singapore

38Instituto Nacional de Salud, Lima, Peru

39Institut Pasteur de Madagascar, Antananarivo, Madagascar

40FIND, Geneva, Switzerland

41University of California, San Diego, USA

42Univ. Lille, CNRS, Inserm, CHU Lille, Institut Pasteur de Lille, U1019 - UMR 9017 - CIIL -Center for Infection and Immunity of Lille, F-59000 Lille, France

43National TB Reference Laboratory, National TB Control Program, Islamabad, Pakistan

44University of Antwerp, Antwerp, Belgium

45University of Edinburgh, Edinburgh, UK

46Stellenbosch University, Cape Town, South Africa

47Wellcome Centre for Infectious Diseases Research in Africa, Cape Town, South Africa

48Francis Crick Institute, London, UK

49Institute of Microbiology, Chinese Academy of Sciences, Beijing, China

## Author contributions

D.W.C, T.E.A.P, S.H, A.L.G.C, A.W.S, T.M.W, P.W.F, D.M.C designed the CRyPTIC study and all contributing laboratories collected samples and provided data. MIC data and genetic information was retrieved and processed by P.W.F, S.H, A.L.G.C, Z.I, M.H and J.K. A.B and K.M.M designed and performed all analyses for this manuscript. A.B and K.M.M wrote the manuscript with feedback from CRyPTIC partners.

## Acknowledgements

We thank Faisal Masood Khanzada and Alamdar Hussain Rizvi (NTRL, Islamabad, Pakistan), Angela Starks and James Posey (Centers for Disease Control and Prevention, Atlanta, USA), and Juan Carlos Toro and Solomon Ghebremichael (Public Health Agency of Sweden, Solna, Sweden), Iñaki Comas and Álvaro Chiner-Oms (Instituto de Biología Integrativa de Sistemas, Valencia, Spain; CIBER en Epidemiología y Salud Pública, Valencia, Spain; Instituto de Biomedicina de Valencia, IBV-CSIC, Valencia, Spain).

## Funding bodies

This work was supported by Wellcome Trust/Newton Fund-MRC Collaborative Award (200205/Z/15/Z); and Bill & Melinda Gates Foundation Trust (OPP1133541). Oxford CRyPTIC consortium members are funded/supported by the National Institute for Health Research (NIHR) Oxford Biomedical Research Centre (BRC), the views expressed are those of the authors and not necessarily those of the NHS, the NIHR or the Department of Health, and the National Institute for Health Research (NIHR) Health Protection Research Unit in Healthcare Associated Infections and Antimicrobial Resistance, a partnership between Public Health England and the University of Oxford, the views expressed are those of the authors and not necessarily those of the NIHR, Public Health England or the Department of Health and Social Care. J.M. is supported by the Wellcome Trust (203919/Z/16/Z). Z.Y. is supported by the National Science and Technology Major Project, China Grant No. 2018ZX10103001. K.M.M. is supported by EMBL’s EIPOD3 programme funded by the European Union’s Horizon 2020 research and innovation programme under Marie Skłodowska Curie Actions.

T.C.R. is funded in part by funding from Unitaid Grant No. 2019-32-FIND MDR. R.S.O. is supported by FAPESP Grant No. 17/16082-7. L.F. received financial support from FAPESP Grant No. 2012/51756-5. B.Z. is supported by the National Natural Science Foundation of China (81991534) and the Beijing Municipal Science & Technology Commission (Z201100005520041). N.T.T.T. is supported by the Wellcome Trust International Intermediate Fellowship (206724/Z/17/Z). G.T. is funded by the Wellcome Trust. R.W. is supported by the South African Medical Research Council.

J.C. is supported by the Rhodes Trust and Stanford Medical Scientist Training Program (T32 GM007365). A.L. is supported by the National Institute for Health Research (NIHR) Health Protection Research Unit in Respiratory Infections at Imperial College London. S.G.L. is supported by the Fonds de Recherche en Santé du Québec.

C.N. is funded by Wellcome Trust Grant No. 203583/Z/16/Z. A.V.R. is supported by Research Foundation Flanders (FWO) under Grant No. G0F8316N (FWO Odysseus).

G.M. was supported by the Wellcome Trust (098316, 214321/Z/18/Z, and 203135/Z/16/Z), and the South African Research Chairs Initiative of the Department of Science and Technology and National Research Foundation (NRF) of South Africa (Grant No. 64787). The funders had no role in the study design, data collection, data analysis, data interpretation, or writing of this report. The opinions, findings and conclusions expressed in this manuscript reflect those of the authors alone. L.G. was supported by the Wellcome Trust (201470/Z/16/Z), the National Institute of Allergy and Infectious Diseases of the National Institutes of Health under award number 1R01AI146338, the GOSH Charity (VC0921) and the GOSH/ICH Biomedical

Research Centre (www.nihr.ac.uk). A.B. is funded by the NDM Prize Studentship from the Oxford Medical Research Council Doctoral Training Partnership and the Nuffield Department of Clinical Medicine. D.J.W. is supported by a Sir Henry Dale Fellowship jointly funded by the Wellcome Trust and the Royal Society (Grant No. 101237/Z/13/B) and by the Robertson Foundation. A.S.W. is an NIHR Senior Investigator. T.M.W. is a Wellcome Trust Clinical Career Development Fellow (214560/Z/18/Z). A.S.L. is supported by the Rhodes Trust. R.J.W. receives funding from the Francis Crick Institute which is supported by Wellcome Trust, (FC0010218), UKRI (FC0010218), and CRUK (FC0010218). T.C. has received grant funding and salary support from US NIH, CDC, USAID and Bill and Melinda Gates Foundation. The computational aspects of this research were supported by the Wellcome Trust Core Award Grant Number 203141/Z/16/Z and the NIHR Oxford BRC. Parts of the work were funded by the German Center of Infection Research (DZIF). The Scottish Mycobacteria Reference Laboratory is funded through National Services Scotland. The Wadsworth Center contributions were supported in part by Cooperative Agreement No. U60OE000103 funded by the Centers for Disease Control and Prevention through the Association of Public Health Laboratories and NIH/NIAID grant AI-117312. Additional support for sequencing and analysis was contributed by the Wadsworth Center Applied Genomic Technologies Core Facility and the Wadsworth Center Bioinformatics Core. SYNLAB Holding Germany GmbH for its direct and indirect support of research activities in the Institute of Microbiology and Laboratory Medicine Gauting. N.R. thanks the Programme National de Lutte contre la Tuberculose de Madagascar.

## Wellcome Trust Open Access

This research was funded in part, by the Wellcome Trust/Newton Fund-MRC Collaborative Award [200205/Z/15/Z]. For the purpose of Open Access, the author has applied a CC BY public copyright licence to any Author Accepted Manuscript version arising from this submission.

This research was funded, in part, by the Wellcome Trust [214321/Z/18/Z, and 203135/Z/16/Z]. For the purpose of open access, the author has applied a CC BY public copyright licence to any Author Accepted Manuscript version arising from this submission.

## Supplemental Information

### S1. Lineages of the *M. tuberculosis* isolates of the compendium

Isolates of the ancient Indo-oceanic lineage/L1 (*n* = 1,150) were mostly contributed by India (*n* = 676 isolates) and Vietnam (*n* = 283 isolates). 85% of the L1 Indian isolates belong to sub-lineages 1.1.2 and 1.2.2 while 66% of the Vietnamese isolates are sub-lineage 1.1.1.1. No L1 isolates were contributed by 10 of the 23 countries in this study with only 2 isolates collected in South America (one in each of Brazil and Peru).

There are 5,598 L2 (East Asian) isolates, making it the second largest group in the dataset. L2 was found most prevalent in Asia and Europe with the largest proportion found in amongst isolates contributed by China (*n* = 722, 64% of isolates) and India (*n* = 1,573, 39% of isolates). Sub-lineages 2.2 and 2.2.7 dominate the L2 isolates (*n* = 1,421 and 1,249 respectively); 2.2 was found mostly amongst Peruvian and South African isolates (*n* = 231 and 161 respectively) apart from those contributed by the Asian countries of Vietnam (*n* = 271), China (*n* = 284) and India (*n* = 272), while 85% of sub-lineage 2.2.7 isolates were contributed by South Africa (*n* = 206), Vietnam (*n* = 164) and India (*n* = 691). 70% of sub-lineage 2.2.1 originated from South Africa (10% of isolates found here) and has recently been associated with more favourable transmission rates (56). Lastly, 86% and 72% of isolates contributed by Kyrgyzstan and Turkmenistan respectively belong to L2 with sub-lineage 2.2.10 dominating (16/24 and 75/86 isolates for both countries respectively). 2.2.10 has been previously described as restricted to Central Asia and this is reflected in the compendium (57).

The majority (1184/1850, 65%) of L3 (East African/Indian) isolates were contributed by India, followed by 19.6% (363/1850) isolates from Pakistan. L3 is typically under-sampled and under-studied in current databases and biobanks in comparison to L2 and L4; the L3 isolates in this study are the largest collected to date in a single study (31).

L4 (Euro-American) is the largest lineage group in the compendium (*n* = 6,572). Isolates donated from Peru dominate; 87% of all L4 isolates are Peruvian with 4.1.2.1 and 4.3.3 being the most prevalent sub-lineages (24% and 22% respectively). There are 34 different L4 sub-lineages in the dataset, making L4 the most diverse in comparison to the other lineage groupings.

There were no L5 isolates found in the compendium and only 6 L6 isolates were identified. Animal-restricted pathogenic mycobacterial isolates are also rare in the compendium; only 16 cases were identified (*n* = 15 *M. bovis* and *n* = 1 *M. caprae*).

---

**Figure S1:**
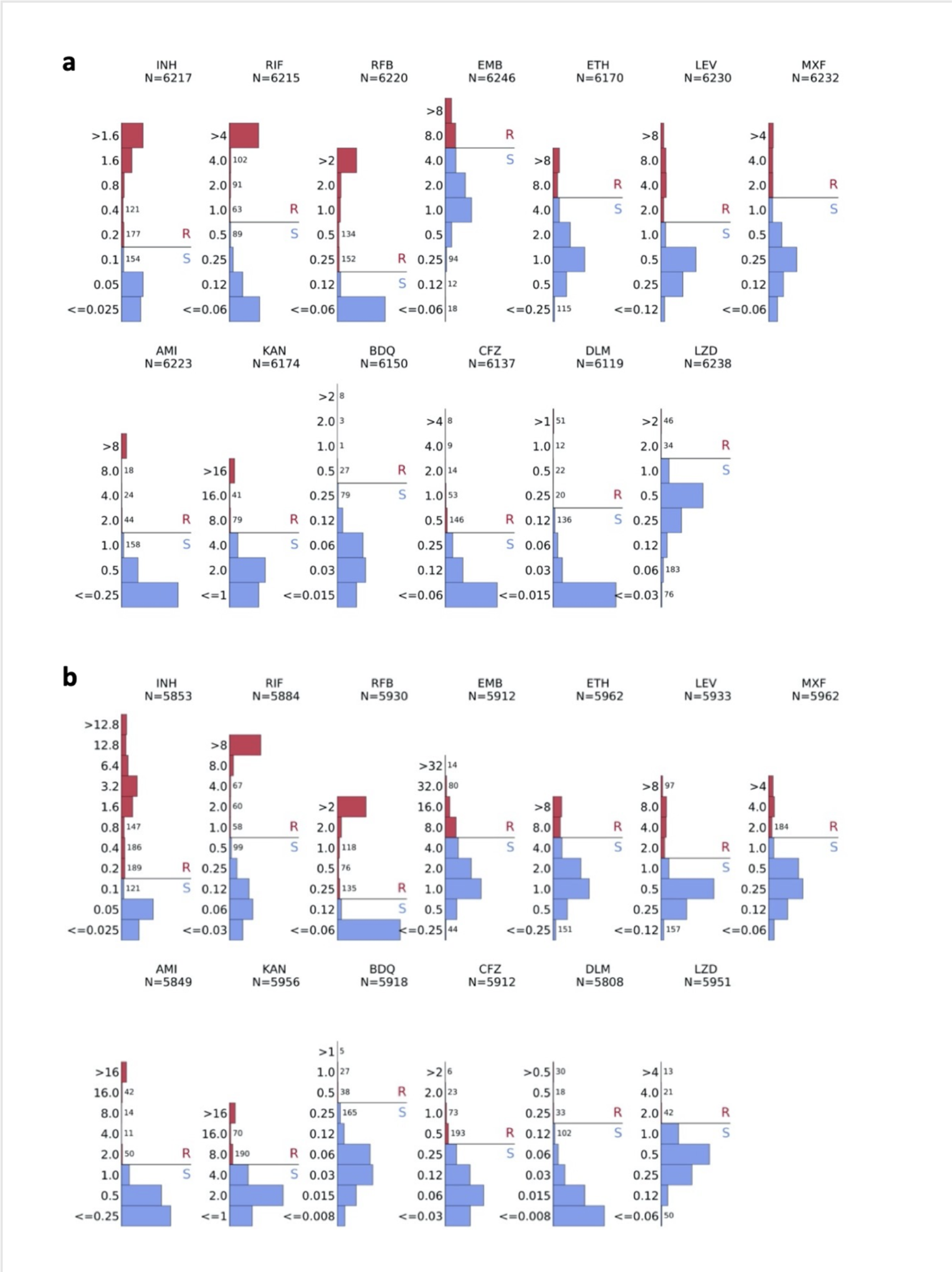
Per drug MIC distributions of isolates plated on CRyPTIC designed variations on the Thermo Fischer Sensititre MYCOTB MIC plate; UKMYC5 (a) and UKMYC6 (b). The solid black line represents the epidemiological resistance cut-off (ECOFF) for each drug as determined by (11). Isolates with an MIC above the cut-off are considered resistant. N denotes the total number of isolates tested on each plate that returned a phenotype for each drug.

**Figure S2:**
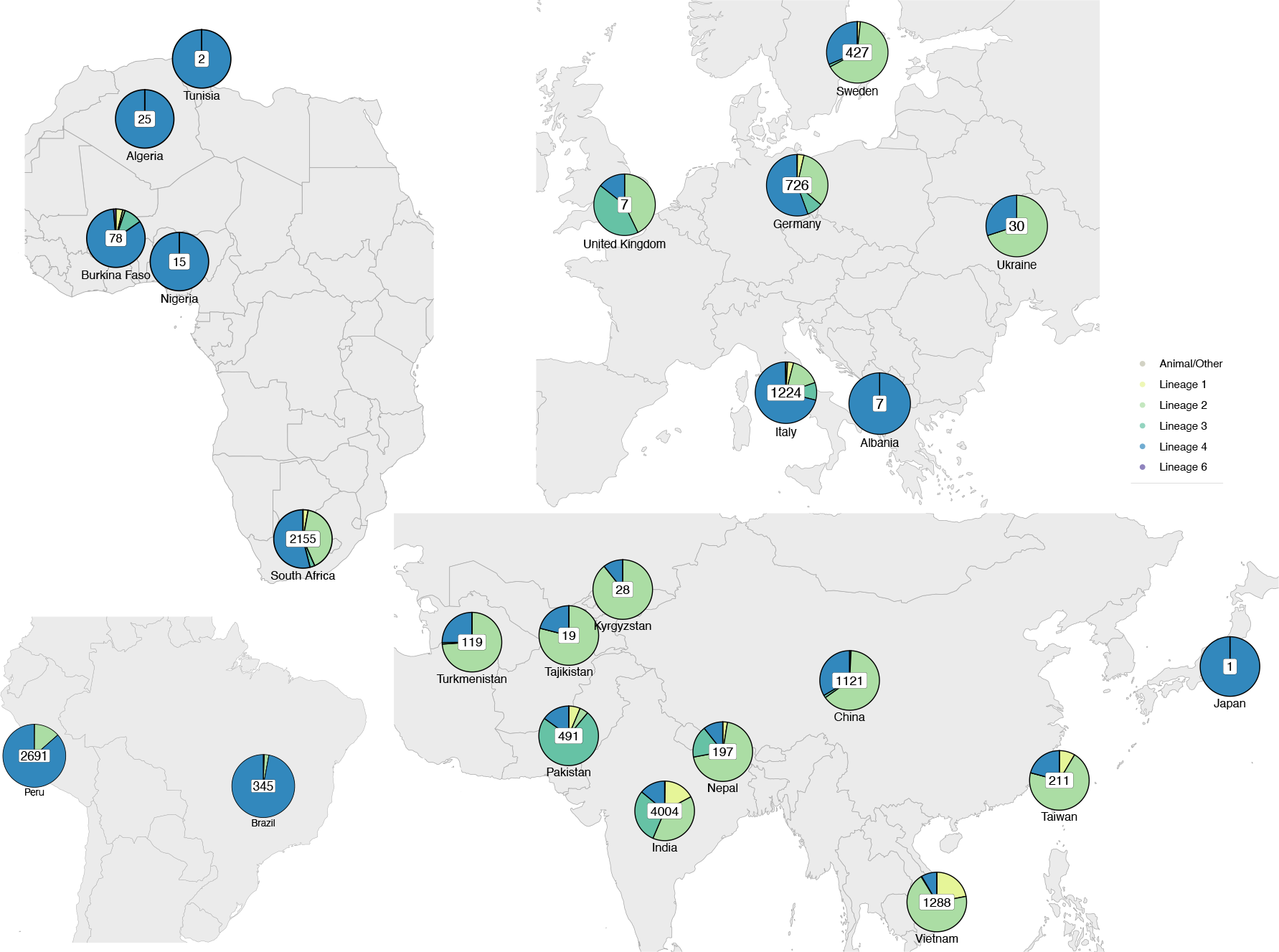
Gographical distribution of CRyPTIC M. tuberculosis clinical isolates. The total number of isolates contributed by each country is depicted, with pie charts representing the proportion of *M. tuberculosis* lineages. Where the origin of an isolate was not known, the collection site identity was assigned (269 isolates in Germany, 17 isolates in India, 6 isolates in Peru, 885 isolates in Italy, 510 isolates in South Africa, 357 isolates in Sweden, 208 isolates in Taiwan, 1 isolate in Brazil and 4 isolates in the UK).

**Figure S3:**
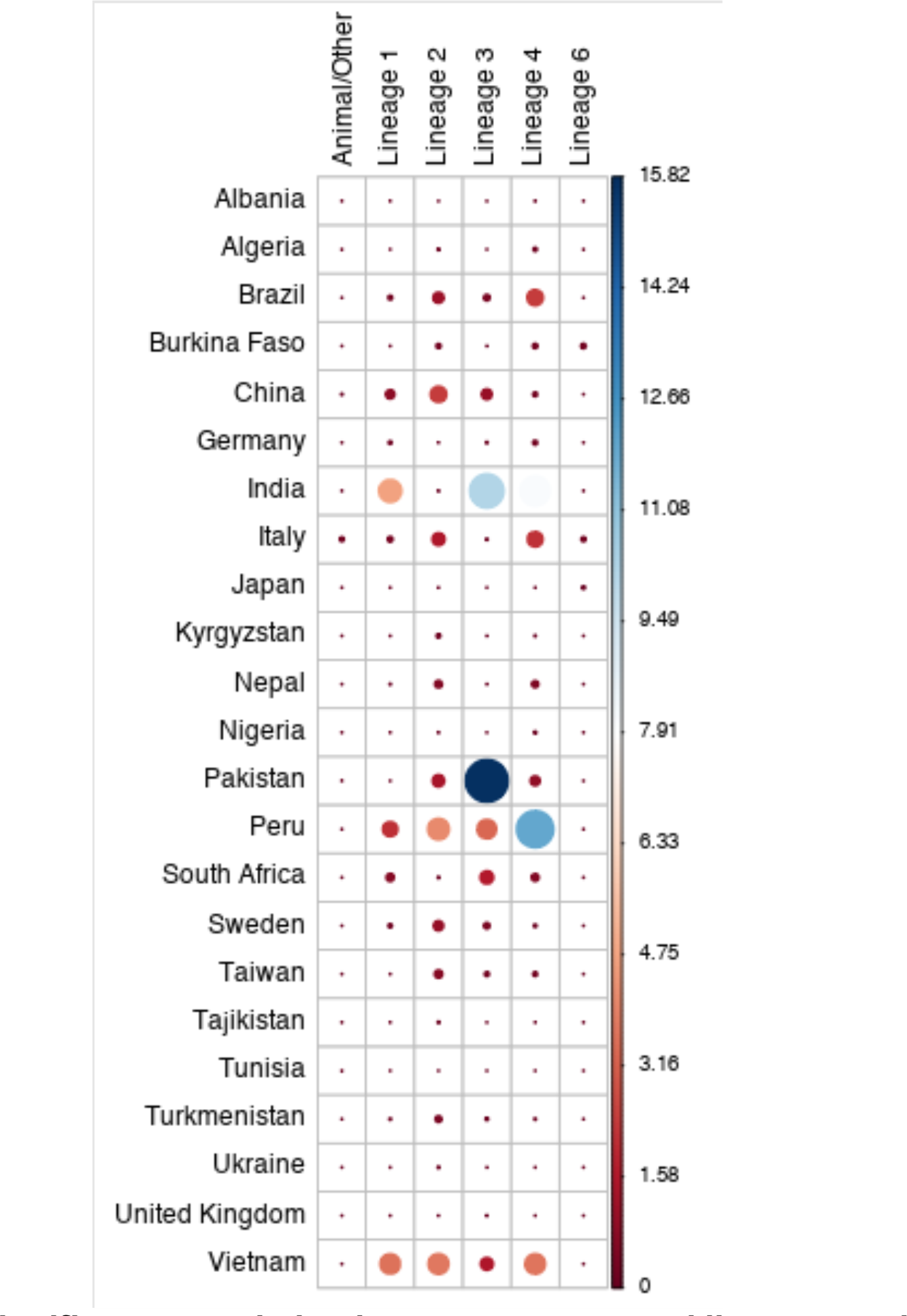
A significant association between country and lineage can be seen in the CRyPTIC data. Pearson’s chi-squared test, X-squared = 7935.2, df = 110, *p* < 2.2e-16. The correlation plot indicates the relative contribution of each row-column pairing to the chi-square test score (%).

**Figure S4:**
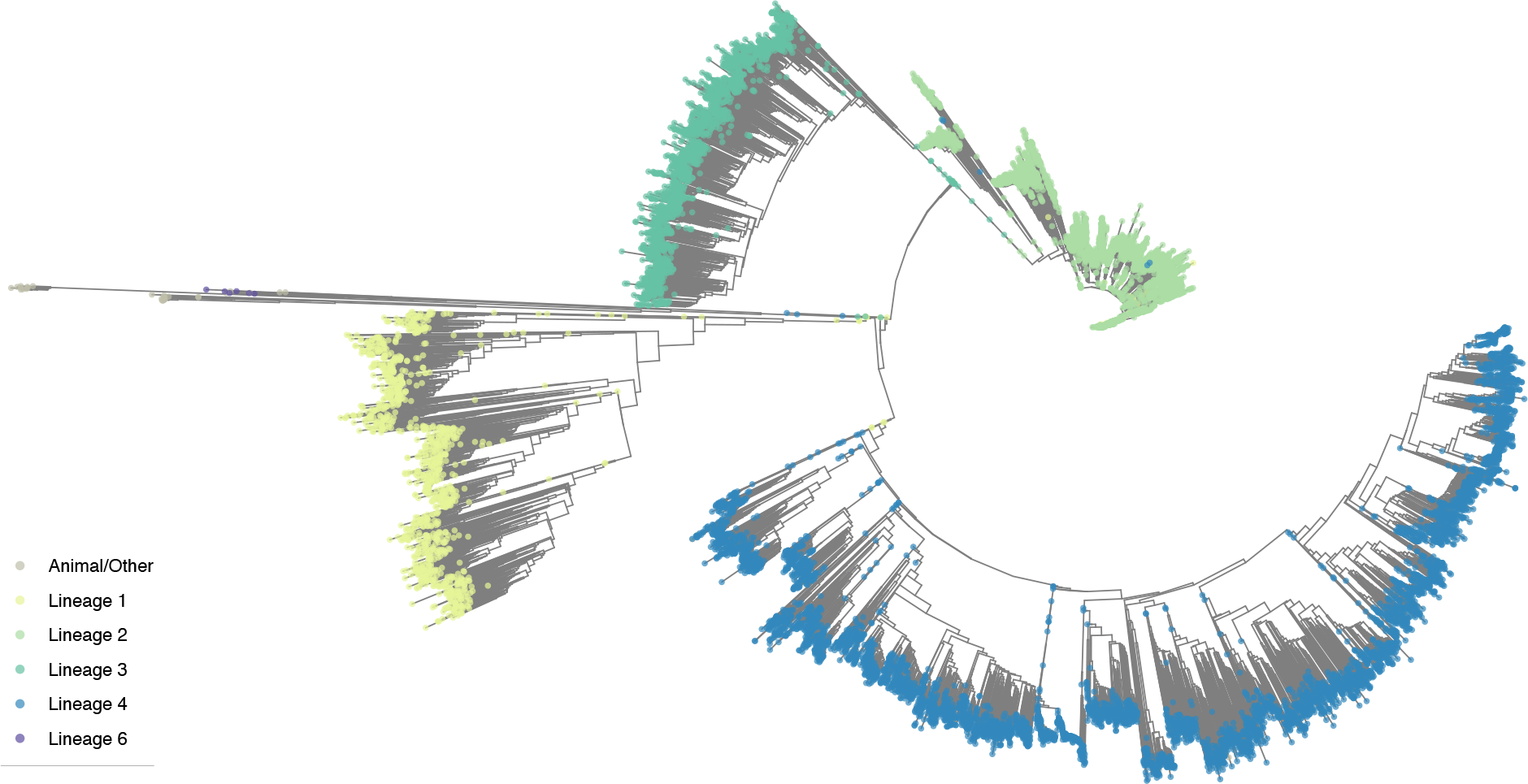
Phylogenetic tree of CRyPTIC M. tuberculosis clinical isolates. A phylogenetic cladogram of 15,211 *M. tuberculosis* clinical isolates. A neighbour-joining tree was constructed from a pairwise distance matrix using *quicktree* (27). Coloured dots at the branch termini represent the lineage assigned to each isolate. “Animal/Other” includes 16 isolates that were assigned the following lineages: *M. caprae* (1)*, M. bovis* (1), along with 17 isolates previously defined as representative for specific sub-lineages (58).

**Figure S6:**
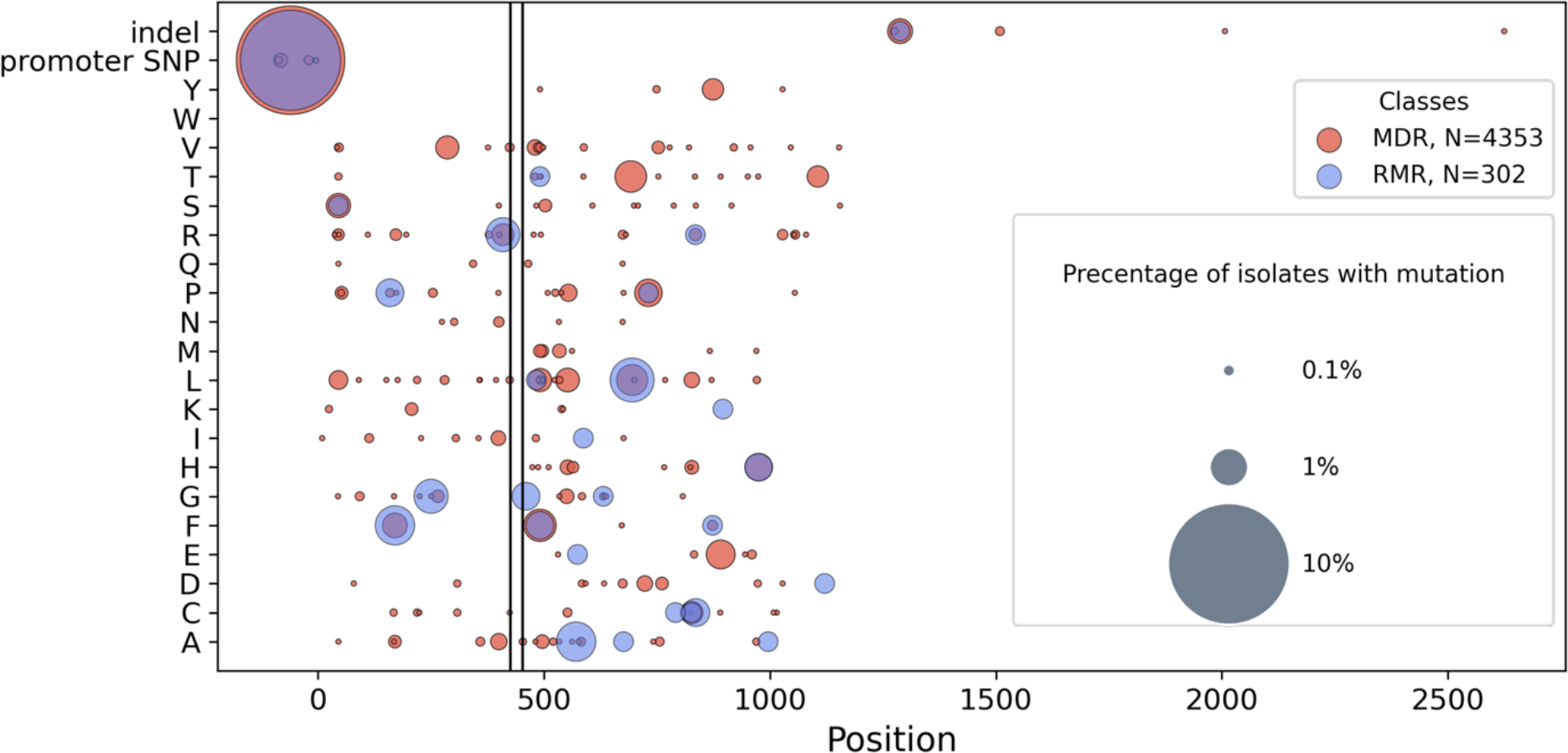
Non-synonymous mutations found outside the RRDR of rpoB in RMR isolates and MDR isolates. Presence of a coloured spot indicates that the mutation was found in RMR/MDR isolates and spot size corresponds to the proportion of RMR or MDR isolates carrying that mutation.

**Table S1:** Epidemiological cut-off values (*ECOFFs) used to binarize MIC measurements into resistant and susceptible.* Isolates with an MIC above the cut-off are considered resistant and those at or below the cut-off as susceptible.

| DRUG | ECOFF |
| --- | --- |
| Isoniazid | 0.1 |
| Rifampicin | 0.5 |
| Rifabutin | 0.12 |
| Ethambutol | 4 |
| Ethionamide | 4 |
| Levofloxacin | 1 |
| Moxifloxacin | 1 |
| Amikacin | 1 |
| Kanamycin | 4 |
| Bedaquiline | 0.25 |
| Clofazimine | 0.25 |
| Delamanid | 0.12 |
| Linezolid | 1 |

**Table S2:**
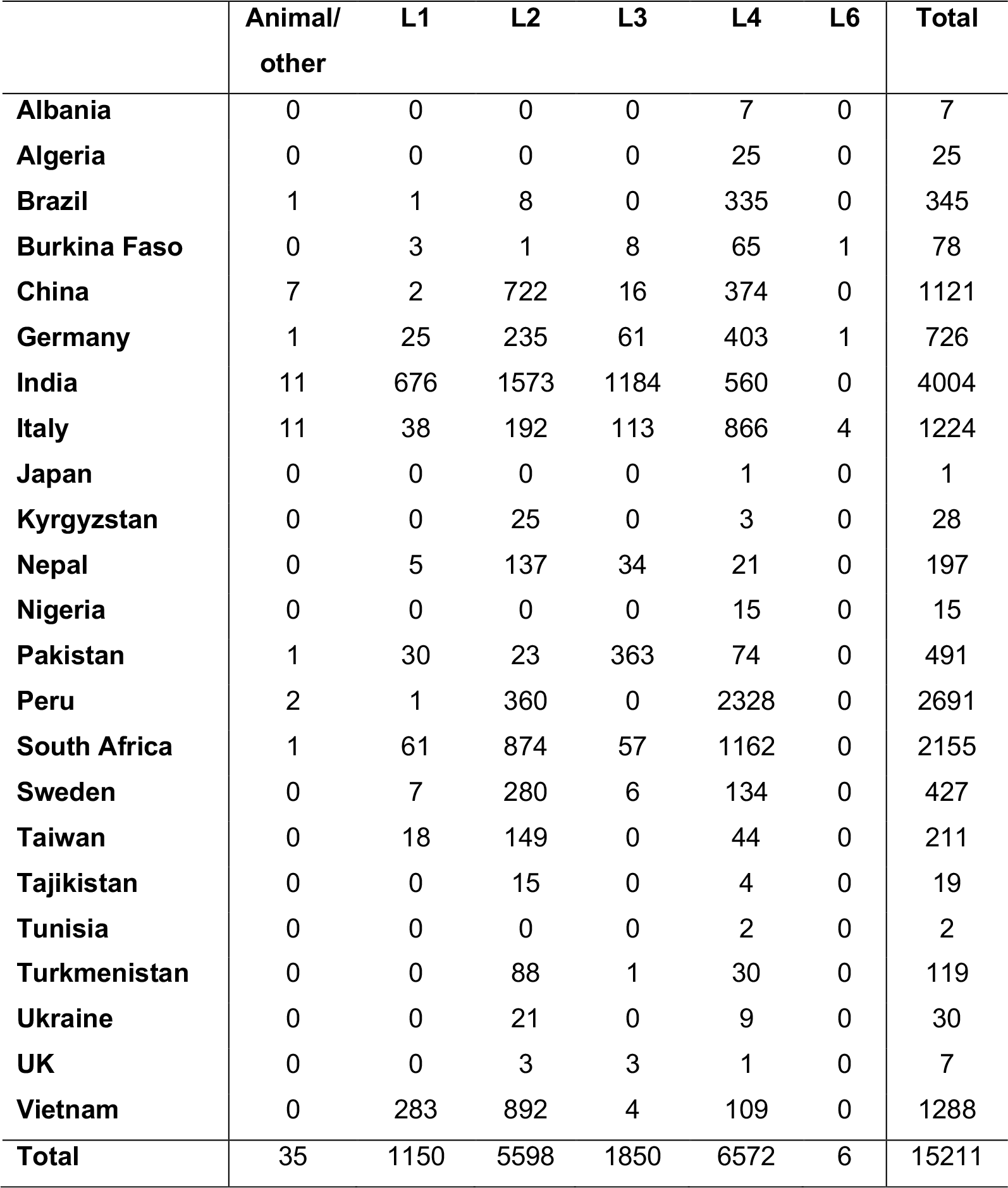
Lineages –v- geographical location of origin/contribution for CRyPTIC isolates.

**Table S3:**
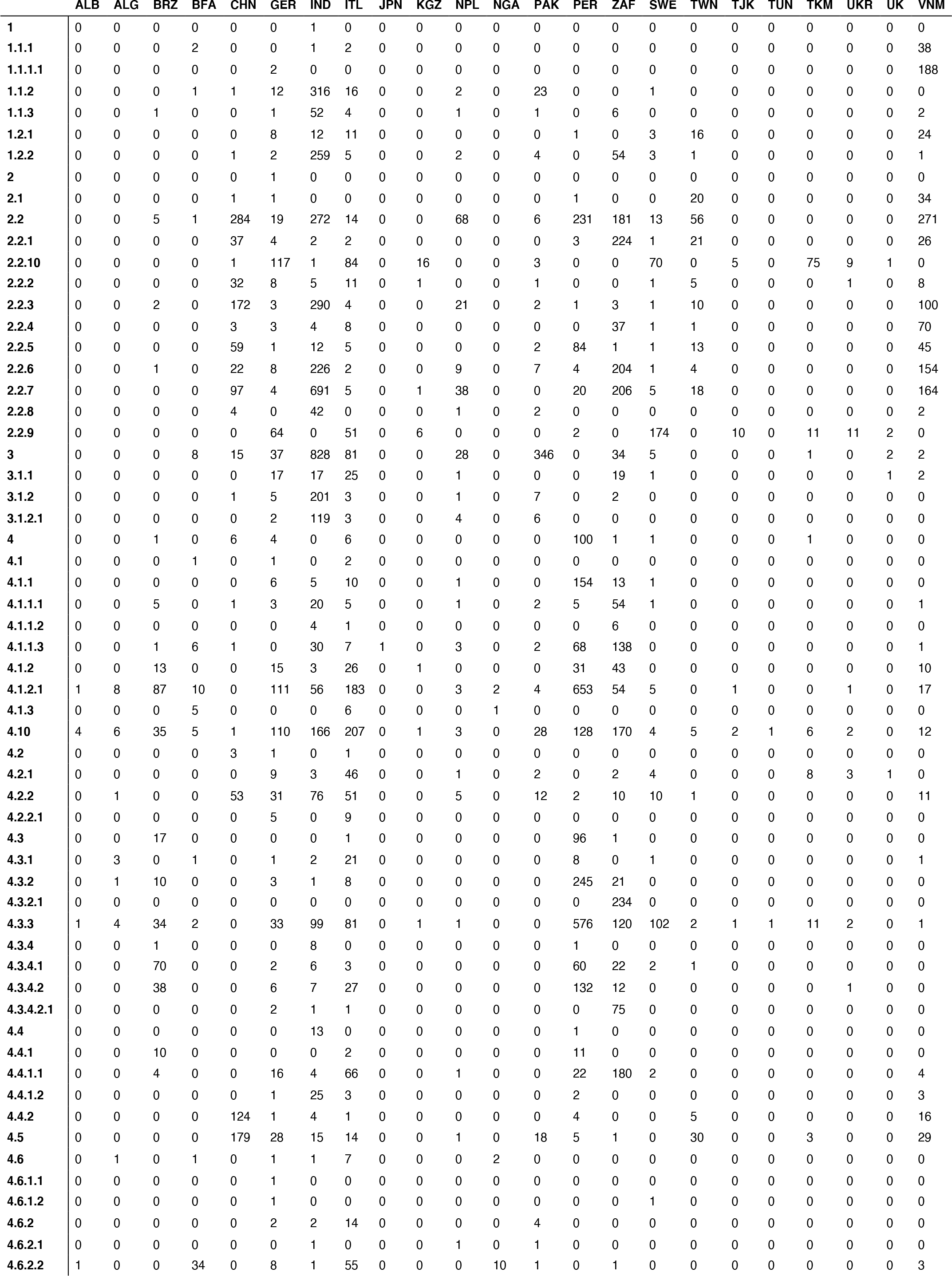
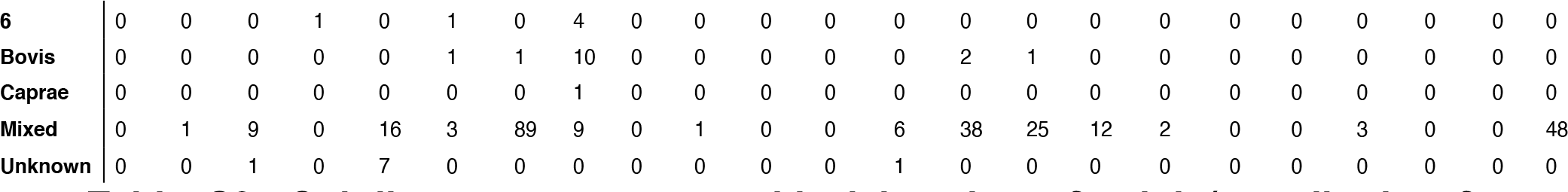
Sub-lineages –v- geographical location of origin/contribution for *CRyPTIC isolates*.

**Table S5:** Sample information for isolates classified as resistant to all 13 CRyPTIC drugs tested. The country of origin is specified using the 3-letter country codes (alpha-3) defined by ISO 3166-1.

| UNIQUEID | COUNTRY OF ORIGIN | LINEAGE |
| --- | --- | --- |
| site.11.subj.XTB-18-224.lab.XTB-18-224.iso.1 | UNKNOWN | Lineage 2 |
| site.10.subj.YA00026182.lab.YA00026182.i so.1 | ZAF | Lineage 4 |

**Table S6:** Co-occurrence of antibiotic resistance in CRyPTIC M. tuberculosis isolates. The probability (%) of an isolate being resistant to Drug 2 (top) if it is resistant to Drug 1 (left). Drug acronyms: INH = isoniazid, RIF = rifampicin, EMB = ethambutol, LEV = levofloxacin, MXF = moxifloxacin, AMI = amikacin, KAN = kanamycin, BDQ = bedaquiline, CFZ = clofazimine, DLM = delamanid, LZD = linezolid, ETH = ethionamide, RFB = rifabutin.

|  | INH | RIF | EMB | LEV | MXF | AMI | KAN | BDQ | CFZ | DLM | LZD | ETH | RFB |
| --- | --- | --- | --- | --- | --- | --- | --- | --- | --- | --- | --- | --- | --- |
| INH | 100.0 | 74.8 | 38.0 | 34.0 | 27.8 | 14.0 | 17.0 | 1.5 | 5.5 | 1.7 | 2.0 | 28.2 | 70.4 |
| RIF | 93.5 | 100.0 | 46.3 | 41.4 | 34.1 | 17.2 | 20.8 | 1.8 | 6.0 | 1.7 | 2.3 | 29.4 | 91.3 |
| EMB | 98.5 | 95.9 | 100.0 | 53.9 | 47.1 | 23.3 | 26.4 | 2.4 | 8.2 | 2.3 | 3.7 | 35.8 | 87.3 |
| LEV | 93.3 | 90.2 | 56.9 | 100.0 | 78.5 | 27.3 | 31.3 | 3.1 | 9.4 | 2.9 | 4.7 | 39.2 | 87.1 |
| MXF | 95.0 | 92.3 | 61.9 | 97.6 | 100.0 | 29.9 | 34.6 | 3.2 | 9.9 | 2.7 | 5.3 | 41.9 | 88.8 |
| AMI | 93.0 | 90.4 | 58.9 | 65.3 | 57.8 | 100.0 | 90.4 | 2.4 | 13.0 | 5.2 | 8.1 | 40.7 | 82.8 |
| KAN | 89.4 | 86.7 | 53.0 | 59.5 | 53.2 | 72.0 | 100.0 | 1.8 | 10.6 | 3.2 | 5.7 | 40.4 | 80.0 |
| BDQ | 79.4 | 77.8 | 49.5 | 60.7 | 49.5 | 20.0 | 18.9 | 100.0 | 52.4 | 12.9 | 14.0 | 34.9 | 76.9 |
| CFZ | 61.9 | 52.8 | 34.9 | 38.2 | 32.1 | 21.8 | 22.5 | 10.6 | 100.0 | 9.4 | 10.2 | 27.0 | 53.7 |
| DLM | 55.2 | 44.4 | 28.2 | 33.2 | 23.9 | 24.4 | 19.1 | 7.1 | 26.3 | 100.0 | 19.3 | 22.1 | 45.4 |
| LZD | 77.6 | 69.9 | 53.2 | 64.7 | 58.7 | 46.7 | 41.2 | 9.8 | 34.2 | 24.1 | 100.0 | 42.6 | 66.7 |
| ETH | 96.5 | 79.4 | 47.0 | 48.7 | 41.9 | 21.1 | 26.2 | 2.2 | 8.3 | 2.4 | 3.9 | 100.0 | 77.1 |
| RFB | 93.3 | 96.8 | 44.5 | 42.1 | 34.4 | 16.6 | 20.3 | 1.9 | 6.4 | 1.9 | 2.3 | 29.9 | 100.0 |

## References

1. WHO. Global Tuberculosis Report 2020. 2021;

2. Shinnick TM, Starks AM, Alexander HL, Castro KG. Evaluation of the Cepheid Xpert MTB/RIF assay. Expert review of molecular diagnostics. 2015 Jan;15(1):9–22.

3. Boehme CC, Nicol MP, Nabeta P, Michael JS, Gotuzzo E, Tahirli R, et al. Feasibility, diagnostic accuracy, and effectiveness of decentralised use of the Xpert MTB/RIF test for diagnosis of tuberculosis and multidrug resistance: a multicentre implementation study. Lancet (London, England). 2011 Apr 30;377(9776):1495–505.

4. Makhado NA, Matabane E, Faccin M, Pinçon C, Jouet A, Boutachkourt F, et al. Outbreak of multidrug-resistant tuberculosis in South Africa undetected by WHO-endorsed commercial tests: an observational study. The Lancet Infectious Diseases. 2018 Dec;18(12):1350–9.

5. Beckert P, Sanchez-Padilla E, Merker M, Dreyer V, Kohl TA, Utpatel C, et al. MDR *M. tuberculosis* outbreak clone in Eswatini missed by Xpert has elevated bedaquiline resistance dated to the pre-treatment era. Genome Medicine. 2020 Dec 25;12(1):104.

6. Sanchez-Padilla E, Merker M, Beckert P, Jochims F, Dlamini T, Kahn P, et al. Detection of Drug-Resistant Tuberculosis by Xpert MTB/RIF in Swaziland. New England Journal of Medicine. 2015 Mar 19;372(12):1181–2.

7. The CRyPTIC Consortium, 100 000 Genomes ProjecT. Prediction of Susceptibility to First-Line Tuberculosis Drugs by DNA Sequencing. New England Journal of Medicine. 2018 Oct 11;379(15):1403–15.

8. Pankhurst LJ, del Ojo Elias C, Votintseva AA, Walker TM, Cole K, Davies J, et al. Rapid, comprehensive, and affordable mycobacterial diagnosis with whole-genome sequencing: a prospective study. The Lancet Respiratory medicine. 2016 Jan;4(1):49–58.

9. Kalokhe AS, Shafiq M, Lee JC, Ray SM, Wang YF, Metchock B, et al. Multidrug-resistant tuberculosis drug susceptibility and molecular diagnostic testing. The American journal of the medical sciences. 2013 Feb;345(2):143–8.

10. Rancoita PM v, Cugnata F, Gibertoni Cruz AL, Borroni E, Hoosdally SJ, Walker TM, et al. Validating a 14-Drug Microtiter Plate Containing Bedaquiline and Delamanid for Large-Scale Research Susceptibility Testing of *Mycobacterium tuberculosis*. Antimicrobial agents and chemotherapy. 2018;62(9).

11. The CRyPTIC Consortium. Epidemiological cutoff values for a 96-well broth microdilution plate for high-throughput research antibiotic susceptibility testing of *M. tuberculosis*. European Respiratory Journal. 2022 Mar 17;2200239.

12. The CRyPTIC Consortium. Quantitative measurement of antibiotic resistance in *Mycobacterium tuberculosis* reveals genetic determinants of resistance and susceptibility in a target gene approach. bioRxiv. 2021;

13. The CRyPTIC Consortium. Predicting Susceptibility to First- and Second-line Tuberculosis Drugs by DNA sequencing and Machine Learning. In preparation.

14. The CRyPTIC Consortium. Genome-wide association studies of global *Mycobacterium tuberculosis* resistance to thirteen antimicrobials in 10,228 genomes. bioRxiv. 2021;

15. Sonnenkalb L, Carter J, Spitaleri A, Iqbal Z, Hunt M, Malone K, et al. Deciphering Bedaquiline and Clofazimine Resistance in Tuberculosis: An Evolutionary Medicine Approach. bioRxiv. 2021 Jan 1;2021.03.19.436148

16. Fowler PW, Gibertoni Cruz AL, Hoosdally SJ, Jarrett L, Borroni E, Chiacchiaretta M, et al. Automated detection of bacterial growth on 96-well plates for high-throughput drug susceptibility testing of *Mycobacterium tuberculosis*. Microbiology. 2018 Dec 1;164(12):1522–30.

17. Fowler PW, Wright C, Spiers-Bowers H, Zhu T, Baeten EML, Hoosdally SW, et al. BashTheBug: a crowd of volunteers reproducibly and accurately measure the minimum inhibitory concentrations of 13 antitubercular drugs from photographs of 96- well broth microdilution plates. bioRxiv. 2021 Jan 1;2021.07.20.453060

18. Hunt M, Letcher B, Malone KM, Nguyen G, Hall MB, Colquhoun RM, et al. Minos: variant adjudication and joint genotyping of cohorts of bacterial genomes. bioRxiv. 2021 Jan 1;2021.09.15.460475.

19. Li H, Durbin R. Fast and accurate short read alignment with Burrows-Wheeler transform. Bioinformatics. 2009 Jul 15;25(14):1754–60.

20. Bolger AM, Lohse M, Usadel B. Trimmomatic: a flexible trimmer for Illumina sequence data. Bioinformatics. 2014 Aug 1;30(15):2114–20.

21. Li H, Handsaker B, Wysoker A, Fennell T, Ruan J, Homer N, et al. The Sequence Alignment/Map format and SAMtools. Bioinformatics (Oxford, England). 2009 Aug 15;25(16):2078–9.

22. Iqbal Z, Caccamo M, Turner I, Flicek P, McVean G. De novo assembly and genotyping of variants using colored de Bruijn graphs. Nature Genetics. 2012 Feb 8;44(2):226–32.

23. Hunt M, Bradley P, Lapierre SG, Heys S, Thomsit M, Hall MB, et al. Antibiotic resistance prediction for *Mycobacterium tuberculosis* from genome sequence data with Mykrobe. Wellcome open research. 2019;4:191.

24. Walker TM, Lalor MK, Broda A, Ortega LS, Morgan M, Parker L, et al. Assessment of *Mycobacterium tuberculosis* transmission in Oxfordshire, UK, 2007–12, with whole pathogen genome sequences: an observational study. The Lancet Respiratory Medicine. 2014 Apr;2(4):285–92.

25. Miotto P, Tessema B, Tagliani E, Chindelevitch L, Starks AM, Emerson C, et al. A standardised method for interpreting the association between mutations and phenotypic drug resistance in *Mycobacterium tuberculosis*. European Respiratory Journal. 2017 Dec;50(6):1701354.

26. Walker TM, Miotto P, Köser CU, Fowler PW, Knaggs J, Iqbal Z, et al. The 2021 WHO Catalogue of *Mycobacterium Tuberculosis* Complex Mutations Associated with Drug Resistance: A New Global Standard for Molecular Diagnostics. SSRN Electronic Journal. 2021;

27. Howe K, Bateman A, Durbin R. QuickTree: building huge Neighbour-Joining trees of protein sequences. Bioinformatics. 2002 Nov 1;18(11):1546–7.

28. Yu G. Using ggtree to Visualize Data on Tree-Like Structures. Current Protocols in Bioinformatics. 2020 Mar 5;69(1).

29. Coscolla M, Gagneux S. Consequences of genomic diversity *in Mycobacterium tuberculosis*. Seminars in Immunology. 2014 Dec;26(6):431–44.

30. Gagneux S. Ecology and evolution of *Mycobacterium tuberculosis*. Nature Reviews Microbiology. 2018 Apr 19;16(4):202–13.

31. Freschi L, Vargas R, Husain A, Kamal SMM, Skrahina A, Tahseen S, et al. Population structure, biogeography and transmissibility of *Mycobacterium tuberculosis*. Nature Communications. 2021 Dec 20;12(1):6099.

32. Ghimire S, Karki S, Maharjan B, Kosterink JGW, Touw DJ, van der Werf TS, et al. Treatment outcomes of patients with MDR-TB in Nepal on a current programmatic standardised regimen: retrospective single-centre study. BMJ Open Respiratory Research. 2020 Aug 12;7(1):e000606.

33. Sharma R, Sharma SK, Singh BK, Mittal A, Kumar P. High degree of fluoroquinolone resistance among pulmonary tuberculosis patients in New Delhi, India. The Indian journal of medical research. 2019 Jan;149(1):62–6.

34. Dijkstra JA, van der Laan T, Akkerman OW, Bolhuis MS, de Lange WCM, Kosterink JGW, et al. In Vitro Susceptibility of *Mycobacterium tuberculosis* to Amikacin, Kanamycin, and Capreomycin. Antimicrobial Agents and Chemotherapy. 2018 Mar;62(3).

35. Maitre T, Petitjean G, Chauffour A, Bernard C, el Helali N, Jarlier V, et al. Are moxifloxacin and levofloxacin equally effective to treat XDR tuberculosis? Journal of Antimicrobial Chemotherapy. 2017 Aug 1;72(8):2326–33.

36. Berrada ZL, Lin S-YG, Rodwell TC, Nguyen D, Schecter GF, Pham L, et al. Rifabutin and rifampin resistance levels and associated rpoB mutations in clinical isolates of *Mycobacterium tuberculosis* complex. Diagnostic Microbiology and Infectious Disease. 2016 Jun;85(2):177–81.

37. Ho J, Jelfs P, Sintchenko V. Fluoroquinolone resistance in non-multidrug-resistant tuberculosis—a surveillance study in New South Wales, Australia, and a review of global resistance rates. International Journal of Infectious Diseases. 2014 Sep;26:149– 53.

38. Singh R, Manjunatha U, Boshoff HIM, Ha YH, Niyomrattanakit P, Ledwidge R, et al. PA-824 Kills Nonreplicating *Mycobacterium tuberculosis* by Intracellular NO Release. Science. 2008 Nov 28;322(5906):1392–5.

39. Hartkoorn RC, Uplekar S, Cole ST. Cross-Resistance between Clofazimine and Bedaquiline through Upregulation of MmpL5 in *Mycobacterium tuberculosis*. Antimicrobial Agents and Chemotherapy. 2014 May;58(5):2979–81.

40. Degiacomi G, Sammartino JC, Sinigiani V, Marra P, Urbani A, Pasca MR. In vitro Study of Bedaquiline Resistance in *Mycobacterium tuberculosis* Multi-Drug Resistant Clinical Isolates. Frontiers in microbiology. 2020;11:559469.

41. Almeida D, Ioerger T, Tyagi S, Li S-Y, Mdluli K, Andries K, et al. Mutations in pepQ Confer Low-Level Resistance to Bedaquiline and Clofazimine in *Mycobacterium tuberculosis*. Antimicrobial Agents and Chemotherapy. 2016 Aug;60(8):4590–9.

42. Nimmo C, Millard J, van Dorp L, Brien K, Moodley S, Wolf A, et al. Population-level emergence of bedaquiline and clofazimine resistance-associated variants among patients with drug-resistant tuberculosis in southern Africa: a phenotypic and phylogenetic analysis. The Lancet Microbe. 2020 Aug;1(4):e165–74.

43. Mvelase NR, Balakrishna Y, Lutchminarain K, Mlisana K. Evolving rifampicin and isoniazid mono-resistance in a high multidrug-resistant and extensively drug-resistant tuberculosis region: a retrospective data analysis. BMJ Open. 2019 Nov 6;9(11):e031663.

44. Villegas L, Otero L, Sterling TR, Huaman MA, van der Stuyft P, Gotuzzo E, et al. Prevalence, Risk Factors, and Treatment Outcomes of Isoniazid- and Rifampicin- Mono-Resistant Pulmonary Tuberculosis in Lima, Peru. PLOS ONE. 2016 Apr 5;11(4):e0152933.

45. Park S, Jo K-W, Lee S do, Kim WS, Shim TS. Treatment outcomes of rifampin- sparing treatment in patients with pulmonary tuberculosis with rifampin-mono- resistance or rifampin adverse events: A retrospective cohort analysis. Respiratory Medicine. 2017 Oct;131:43–8.

46. Salaam-Dreyer Z, Streicher EM, Sirgel FA, Menardo F, Borrell S, Reinhard M, et al. Rifampicin-Monoresistant Tuberculosis Is Not the Same as Multidrug-Resistant Tuberculosis: a Descriptive Study from Khayelitsha, South Africa. Antimicrobial Agents and Chemotherapy. 2021 Oct 18;65(11).

47. Pang Y, Lu J, Wang Y, Song Y, Wang S, Zhao Y. Study of the Rifampin Monoresistance Mechanism in *Mycobacterium tuberculosis*. Antimicrobial Agents and Chemotherapy. 2013 Feb;57(2):893–900.

48. Kigozi E, Kasule GW, Musisi K, Lukoye D, Kyobe S, Katabazi FA, et al. Prevalence and patterns of rifampicin and isoniazid resistance conferring mutations in *Mycobacterium tuberculosis* isolates from Uganda. PLOS ONE. 2018 May 30;13(5):e0198091.

49. Dean AS, Zignol M, Cabibbe AM, Falzon D, Glaziou P, Cirillo DM, et al. Prevalence and genetic profiles of isoniazid resistance in tuberculosis patients: A multicountry analysis of cross-sectional data. PLOS Medicine. 2020 Jan 21;17(1):e1003008.

50. Migliori GB, Langendam MW, D’Ambrosio L, Centis R, Blasi F, Huitric E, et al. Protecting the tuberculosis drug pipeline: stating the case for the rational use of fluoroquinolones. European Respiratory Journal. 2012 Oct;40(4):814–22.

51. Heysell SK, Ahmed S, Rahman MdT, Akhanda MdW, Gleason AT, Ebers A, et al. Hearing loss with kanamycin treatment for multidrug-resistant tuberculosis in Bangladesh. European Respiratory Journal. 2018 Mar;51(3):1701778.

52. WHO. Update on the use of nucleic acid amplification tests to detect TB and drug- resistant TB: rapid communication. 2021;

53. Olaru ID, Heyckendorf J, Andres S, Kalsdorf B, Lange C. Bedaquiline-based treatment regimen for multidrug-resistant tuberculosis. European Respiratory Journal. 2017 May 21;49(5):1700742.

54. WHO. World Health Organization treatment guidelines for drug-resistant tuberculosis, 2016 update. 2016;

55. Huitric E, Verhasselt P, Koul A, Andries K, Hoffner S, Andersson DI. Rates and Mechanisms of Resistance Development in *Mycobacterium tuberculosis* to a Novel Diarylquinoline ATP Synthase Inhibitor. Antimicrobial Agents and Chemotherapy. 2010 Mar;54(3):1022–8.

56. Holt KE, McAdam P, Thai PVK, Thuong NTT, Ha DTM, Lan NN, et al. Frequent transmission of the *Mycobacterium tuberculosis* Beijing lineage and positive selection for the EsxW Beijing variant in Vietnam. Nature Genetics. 2018 Jun 21;50(6):849–56.

57. Chiner-Oms Á, Comas I. Large genomics datasets shed light on the evolution of the Mycobacterium tuberculosis complex. Infection, Genetics and Evolution. 2019 Aug;72:10–5.

58. Borrell S, Trauner A, Brites D, Rigouts L, Loiseau C, Coscolla M, et al. Reference set of *Mycobacterium tuberculosis* clinical strains: A tool for research and product development. PLOS ONE. 2019 Mar 25;14(3):e0214088.

